# Establishing insect community composition using metabarcoding of soil samples, and preservative ethanol and homogenate from Malaise trap catches: surprising inconsistencies between methods

**DOI:** 10.1101/597302

**Authors:** Daniel Marquina, Rodrigo Esparza-Salas, Tomas Roslin, Fredrik Ronquist

## Abstract

DNA metabarcoding allows the analysis of insect communities faster and more efficiently than ever before. However, metabarcoding can be conducted through several alternative approaches, and the consistency of results across methods has rarely been studied. We compare the results obtained by DNA metabarcoding of the same communities using two different markers – COI and 16S – and three different sampling methods – homogenized Malaise trap samples (homogenate), preservative ethanol from the same samples, and soil samples. Our results indicate that COI and 16S offer partly complementary information on Malaise trap samples, with each marker detecting a significant number of species not detected by the other. Different sampling methods offer highly divergent estimates of community composition. The community recovered from preservative ethanol of Malaise trap samples is quite distinct from that recovered from homogenate. Small and weakly sclerotized insects tend to be overrepresented in ethanol, with some exceptions that could be related to taxon-specific traits. For soil samples, highly degenerate COI primers pick up large amounts of non-target DNA and only 16S provides adequate analyses of insect diversity. However, even with 16S, very little overlap in MOTU content was found between the trap and the soil samples. Our results demonstrate that no metabarcoding approach is all-comprehensive in itself. For instance, DNA extraction from preservative ethanol is not a valid replacement for destructive bulk extraction but a complement. In future metabarcoding studies, both should ideally be used together to achieve comprehensive representation of the target community.

## Introduction

The spectacular diversity of insects, and the shortage of adequate expertise and resources for taxonomic identification, has long hindered studies of the structure and functioning of the insect component of ecosystems. However, the advent of DNA metabarcoding is about to change this. These new methods promise to make it feasible to analyze the composition and structure of entire insect communities in detail, opening up research questions in community ecology that have simply not been possible to tackle before (Gibson *et al*., 2015).

As exciting as these prospects may be, the methodology of DNA metabarcoding of insect communities is still immature. Several different approaches have been tried but it is still unclear how consistent results are across methods and what protocol is optimal for a particular study (but see Majaneva, Diserud, Eagle, Hajibabaei, & Ekrem, 2018a). For instance, there is still debate over the optimal marker for DNA metabarcoding of insect communities, and to what extent combinations of markers can be used to improve the resolution and accuracy of the analyses (Andújar, Arribas, Yu, Vogler, & Emmerson, 2018; Clarke, Soubrier, Weyrich, & Cooper, 2014; Gibson *et al*., 2014). The efficiency and biases of different sampling and analysis protocols are also poorly understood. For instance, insect communities can be studied using metabarcoding of the preservative ethanol from Malaise trap samples, from homogenized tissue from such samples, or from environmental DNA (eDNA) in soil samples, but we lack an understanding of consistency of results across these methods. This paper is devoted to exploring such methodological issues associated with metabarcoding studies of insect communities.

Multi-marker metabarcoding approaches were initially adopted, and still are, as a solution to obtain biodiversity information from several domains of life in a particular habitat, combining the most used markers for each of the taxa: 16S for prokaryotes, 18S for eukaryotes, COI for animals and rcbl or matK for plants (Drummond *et al.*, 2015; Gibson *et al.*, 2014; Krehenwinkel *et al.*, 2018). In other cases, two markers are used to provide complementary data on the same taxonomic fraction of the sample. In such cases, it is common to use a conserved marker with high taxonomic coverage (such as 18S) to ensure the detection of all the main groups of the target taxon, and a more variable marker (such as COI) providing information at higher taxonomic resolution but at the cost of some degree of amplification bias (Cowart *et al.*, 2015; Tang *et al.*, 2012). This approach of combining multiple markers has the advantage that the two markers complement each other, with one marker ameliorating potential biases in amplification of a specific taxon by the other marker (Freeland, 2017; Wangensteen, Palacín, Guardiola, & Turon, 2018). Yet, the lack of resolution of the conserved marker decreases its effectiveness. In response, some authors have advocated the use of a complementary marker that is more conserved than the high-resolution marker (typically COI), but more variable than the nuclear rRNA genes.

An ideal complementary marker should be easily amplified through the different subtaxa of the target group, while providing enough taxonomic resolution to discriminate between related species. The mitochondrial rRNA gene 16S has been offered as such a marker (Clarke *et al*., 2014; Deagle, Jarman, Coissac, Pompanon, & Taberlet, 2014; Marquina, Andersson, & Ronquist, 2018). To our knowledge, only a few studies have tested the potential of using 16S as a replacement for COI in insects in vitro (Clarke *et al.*, 2014; Elbrecht *et al.*, 2016; Epp *et al.*, 2012), or as a complement (Alberdi, Aizpurua, Gilbert, & Bohmann, 2018), and only one study made use of both markers simultaneously in resolving ecological rather than methodological questions (Kaunisto, Roslin, Sääksjärvi, & Vesterinen, 2017). These studies have reported that the performance of 16S is considerably poorer compared to COI (Alberdi *et al*., 2018; Elbrecht *et al*., 2016), but this discrepancy may be attributed to the fragment of the 16S gene amplified being very short (160–190 bp approx.), and to the fact that the reference library of the 16S gene is substantially smaller than that of COI (Andújar *et al*., 2018; Deagle *et al*., 2014). Using longer fragments of the 16S gene, the taxonomic resolution increases to levels comparable to COI (Clarke *et al*., 2014, Marquina *et al*., 2018). Wilson, Brandon-Mong, Gan & Sing (2018) compared 16S metabarcoding with 16S- and COI-oriented metagenomics and metatranscriptomics, obtaining satisfactory results with respect to taxa detection, but not with respect to abundance measurement. Nevertheless, as they did not include COI metabarcoding, the comparison is difficult.

Malaise traps have been widely used in morphology-based biodiversity surveys (*e.g.* Global Malaise Trap Program (GMP, http://biodiversitygenomics.net/projects/gmp/)). More recently, catches from such traps have been among the first types of bulk samples subjected to DNA metabarcoding analysis (Yu *et al.*, 2012; Gibson *et al.*, 2014, Shokralla *et al.*, 2015), offering promising vistas for automated, large-scale biomonitoring of complex communities (see Porter & Hajibabaei, 2018). Yet, the optimization of protocols for metabarcoding of Malaise trap samples and other terrestrial arthropod samples (*e.g.* canopy fogging) has lagged behind the methods for other types of mixed samples, such as freshwater or marine benthos, or soil. Only a few studies have focused exclusively on methodological improvements of samples of this type (Krehenwinkel *et al.*, 2017; Krehenwinkel *et al.*, 2018; Morinière *et al.*, 2016, Wilson *et al*., 2018). Much of the work done on metabarcoding of freshwater invertebrates can be applied to terrestrial bulk samples. Such transfer of methods may include, for instance, the use of the preservative ethanol to extract DNA without destroying the morphology of the organisms. Nevertheless, insect communities found in freshwater benthos and Malaise traps have different characteristics – with freshwater samples dominated by larval stages, and by low species richness and abundances, while Malaise trap catches are dominated by adult insects, and by high species richness and abundances. As a consequence, the transfer of techniques between these types of samples demands specific testing and optimizing.

An alternative to analyzing DNA from trap catches is to use traces of DNA present in the environment. When successful, such environmental DNA (eDNA) samples — *e.g.*, soil or water — may reveal a large fraction of taxa present in the region, and serve as an efficient indicator of local biodiversity (Bohmann *et al*., 2014; Creer *et al*., 2014; Deiner *et al*., 2017). However, most of the template DNA in environmental samples consists of highly degraded extracellular DNA from decomposed tissues, cells and excreta of organisms. The low quality of the template DNA makes extraction and amplification extra challenging, and primers that are successful for metabarcoding of trap samples may not work for environmental samples. Importantly, eDNA analysis is focused on recovering traces of total biodiversity of the medium the sample was taken from. This can result in a case where the PCR or extraction step are saturated with off-target DNA, thus yielding few sequences of the intended taxon. Highly degenerate primers worsen this situation, especially when such high degeneracy is needed to amplify the full diversity of the target taxon (Horton *et al.*, 2017; Macher *et al.*, 2017). Even when less degenerate primers are used, like those for 18S, DNA amplified from non-desired organisms can still form half of the sequencing output (Yang *et al.*, 2014). Studies comparing the results of metabarcoding of eDNA and trap samples have been scarce, and mainly focused on freshwater samples (Deiner, Fronhofer, Mächler, Walser, & Altermatt 2016; Macher *et al.*, 2017), or soil (Horton, Kershner, & Blackwood 2017). To our knowledge, only one study has combined both types of sample with an ecological aim rather than a methodological one (Yang *et al.*, 2014).

Analyses of complex community samples have oftentimes been based on extraction from the homogenized tissue present in the sample (Aylagas, Mendibil, Borja, & Rodríguez-Ezpeleta, 2016; Emilson *et al*., 2017; Kocher *et al*., 2017). As an alternative approach, Hajibabaei, Spall, Shokralla, & van Konynenburg (2012) and others (Erdozain *et al*., 2019; Zizka, Leese, Peinert, & Geiger, 2018) have suggested extraction from the preservative medium – *i.e.* the ethanol in which the samples are stored. If successful, such an approach would reduce the time of processing and handling, which would decrease the risk of cross-contamination. It would also leave the insects intact. This would not only allow further taxonomic work, but might also allow investigations of the microbiome, parasites/parasitoids or diet of individual specimens already metabarcoded. However, metabarcoding of preservative fluid may well differ from metabarcoding of homogenized tissue. For instance, small insects with high surface/volume ratio might be expected to be represented by proportionately more DNA in the preservative fluid than in the homogenized tissue, since the leakage of DNA to the preservative ethanol is dependent on body surface. When bulk samples of tissue are homogenized, large-sized organisms will contribute much more DNA to the pool than small ones, biasing the proportion of reads per specimen in the opposite direction (Bista *et al*., 2017; Elbrecht, Peinert, & Leese, 2017; Krehenwinkel *et al.*, 2017).

In this study, we extend a previous *in silico* study of markers for insect metabarcoding (Marquina *et al*., 2018) by testing the optimal 16S and COI markers identified there in three different empirical settings. Targeting Malaise trap catches from three different sites at four different time points, we analyse homogenized insects and preservative ethanol from the same samples, and environmental DNA from soil samples taken at the same sites. This allows us to examine to what extent COI and 16S results overlap or complement each other, and to what extent the sample type affects the performance of the two markers. It also allows us to explore the degree of consistency in metabarcoding results across two different markers and three different protocols of sampling and to use these results in analyzing the composition of the focal insect communities.

## Materials and methods

### Study area and sampling

To obtain complex environmental samples of insect communities, we targeted the Nacka Nature Reserve at the edge of the metropolitan area of Stockholm (Sweden). Three sites (detailed in Table S1) were chosen to maximize diversity of habitats. The insect community was sampled at four different time points (T1, T2, T3 and T4, corresponding to July 4^th^, August 1^st^, August 29^th^ and September 26^th^ of 2016 respectively) using a Malaise trap run for a duration of one week and a soil sample. (Figure 1). Each Malaise trap was fitted with a collecting jar filled with 95 % ethanol. Each soil sample consisted of three subsamples taken at random points within a 20 m radius centred at the Malaise trap. Each subsample was taken using a cylindrical core extractor, 30 cm long × 10 cm diameter, dug down to the mineral soil or to its maximum length. Both Malaise trap and soil samples were frozen at −25 °C and −80 °C respectively between 2–3 h after collection, and they were stored for approximately 6 months before analysis. Unique bottles were used for each Malaise trap sample. The core extractor was subjected to a bleach bath to avoid cross-contamination between sites, and soil samples were stored in individual zip-lock bags.

**Figure 1.**
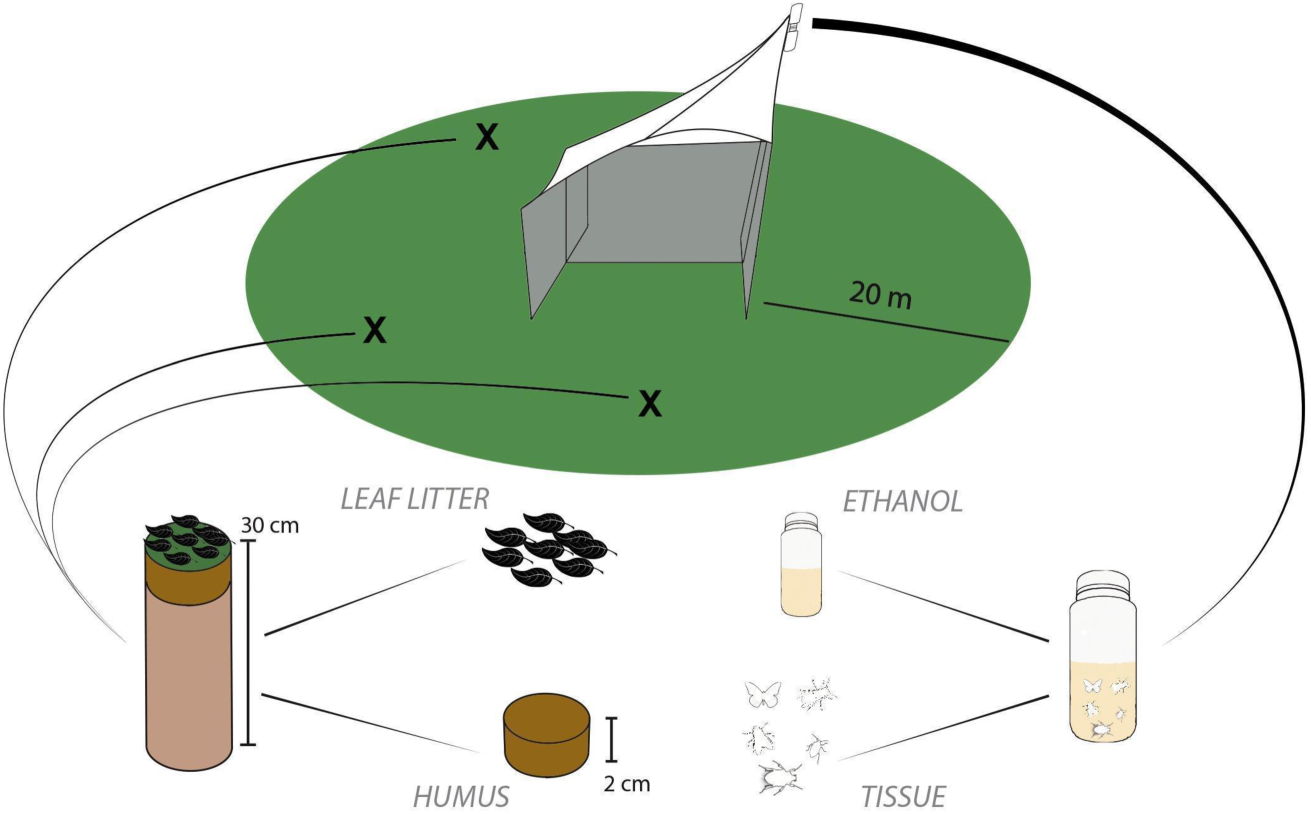
Overview of the sampling process and sample typification. At each time point one Malaise trap and three soil cores were taken. The preservative ethanol of the catch was separated and filtered (Ethanol), and the insects were homogenized (Tissue). Each of the three soil cores were separated in leaf litter (Leaf litter) and the two first centimetres of soil (Humus). The three samples for each time point (3 Humus and 3 Leaf litter) were ground and pooled together.

### DNA extraction

Once in the laboratory, the leaf litter (L hereafter) was isolated from each soil subsample, as well as the first two centimetres of soil (humus or H hereafter). Then, 50 mL of the humus and leaf litter of each subsample were pooled together by sample, respectively, after which each L and H sample was mixed, manually grinded in a mortar after freezing with liquid N_2_ and mixed again, resulting in a total of 24 samples: 2 layers (humus and leaf litter) times 3 sites times 4 time points. Every utensil used to manipulate the soil samples (mortar, shovel, etc.) was washed with soap, bathed in bleach, thoroughly rinsed and UV-irradiated between samples. Gloves and filter paper were discarded after handling each sample and the bench was cleaned with bleach. Subsequently, 0.4 g of each sample were used for DNA extraction using Nucleospin Soil Kit (Macherey-Nagel, Düren, Germany) following the manufacturer’s protocol.

The Malaise trap samples were handled in a special room of the laboratory dedicated to environmental DNA. This space undergoes frequent cleaning and decontamination and all personnel is required to wear special lab coats reserved for this lab. The ethanol of the Malaise trap samples was poured through a sieve of 0.6 mm pore size to retain small individuals and body parts. The ethanol was then filtered using Durapore membrane filters (Merk, Darmstadt, Germany) of 0.45 µm pore size, and discarded. The Durapore filters with the DNA in the preservative ethanol were immediately stored in a 2 mL tube with 400 µL of lysis buffer and stored at −20 °C until extraction, while the insects were left to dry on filter paper. The dried insects were homogenized after freezing with liquid N_2_ and stored in tubes with 400, 800 or 1200 µL of lysis buffer depending on the resulting volume of powder after homogenization. All non-disposable utensils and the working benches were washed, bleached and UV irradiated between samples using the same procedures as for the soil samples. Both the filter membranes (henceforth referred to as ‘ethanol-DNA’ or E) and the homogenized insects (‘tissue-DNA’ or T hereafter) were incubated with proteinase K at 56 °C for 24 h, resulting in a total of 24 samples: 2 extracts (ethanol and tissue) times 3 traps times 4 time points. For each sample, 225 µL were used for DNA extraction with a KingFisher Cell & Tissue DNA kit (Thermo Fisher Scientific, Waltham, USA) on a KingFisher Duo instrument (Thermo Fisher Scientific).

### PCR and sequencing

Two mitochondrial markers were selected for PCR amplification based on the findings of Marquina *et al.*, (2018). The COI marker was amplified using the primer pair BF2–BR1 (Elbrecht, & Leese 2017; see also Marquina et al. 2018), resulting in an amplicon of approx. 320 bp. The 16S marker was amplified using the primer pair Chiar16SF–Chiar16SR (Marquina *et al.*, 2018), resulting in an amplicon of approx. 350 bp. Primers were tagged with a unique sequence of 8 nucleotides at the 5’ end (Binladen *et al*., 2007), and used in different combination of forward and reverse tags for multiplexing samples in a single sequencing run. The PCR mix consisted of one Illustra Hot Start Mix RTG bead (GE Healthcare Life Sciences, Freiburg, Germany), 10 pmoles of each primer, 2 µL of DNA template and 21 µL of biology-grade water (final volume: 25 µL). The temperature protocol consisted of an initial phase of denaturalization and Taq-activation for 5 min at 95 °C, followed by 40 cycles of 95 °C for 30 s, annealing phase at 48 / 50 °C (for COI and 16S respectively) for 45 s and extension phase at 68 °C for 45 s, and a final extension phase at 72 °C for 10 minutes. Blanks were prepared with the same volumes but substituting the DNA with water; the blanks were discarded after checking there was no amplification product in an agarose gel. All PCR runs were duplicated and sample replicates were pooled before library preparation. Final PCR products were pooled to equimolar concentrations and purified, and libraries prepared with TrueSeq PCR-free kit (Illumina, San Diego, USA). Desired DNA fragments with adapters were cut off from an agarose gel, purified using QIAquick gel extraction kit (Qiagen, Hilden, Germany), pooled again to equimolar concentrations and sequenced on an Illumina MiSeq, using v3 chemistry and a 2 × 300 bp paired-end run, at SciLifeLab (Stockholm, Sweden).

### Bioinformatic analysis

Bioinformatic analysis was conducted using a combination of the OBITools package (Boyer *et al.*, 2016), other programs detailed below, custom scripts (available at https://github.com/metagusano/metabarcoding_scripts) and scripts from https://github.com/metabarpark/R_scripts_metabarpark. Read quality was checked using FastQC (Andrews, 2010), and the reads were trimmed after the position in which the average phred score was lower than 28. Paired-end reads with a phred score lower than 30 were discarded. The remaining reads were demultiplexed using the unique sample tags and the primers removed with *ngsfilter*, and selected based on length. For COI, only reads with a length in the interval 310–330 bp and with no ambiguous bases were retained, while the 16S reads were separated in two categories: 16S–‘long’ (315–375 bp and with no ambiguous bases) and 16S–‘short’ (255–305 bp and with no ambiguous bases). These lengths were selected based both on previous results (Marquina *et al.*, 2018, and *in silico* PCR of arachnid sequences (data not shown)) and on the observed length distribution of the reads, suggesting different amplicon lengths for insects and arachnids. The length constraint used for 16S needs to be relaxed because of the increased frequency of indels in rRNA compared to protein-coding genes. From this point on, the pipeline was conducted in parallel for the three markers (COI, 16S–‘long’ and 16S–‘short’). Reads were de-replicated with *obiuniq* and chimera removal was conducted using VSEARCH v2.7.1 (Rognes, Fouri, Nichols, Quince, & Mahé 2016; https://github.com/torognes/vsearch), and the remaining sequences were clustered into molecular operational taxonomic units (MOTUs) using SWARM v2.1.13 (Mahé, Rognes, Quince, de Vargas, & Dunthorn 2015; https://github.com/torognes/swarm). Different distance thresholds were tested for the three markers based on previous knowledge of the barcoding gap for each one (Marquina *et al.*, 2018), and the optimal threshold distance, *d*, was taken to be the value at which the decrease in number of MOTUs reached a stable phase when increasing distance thresholds. The optimal thresholds found using this procedure were: *d* = 4 for 16S–‘short’, *d* = 5 for 16S–‘long’ and *d* = 9 for COI (Figure S1). Resulting MOTU tables were curated with LULU v0.1.0 (Frøslev *et al.*, 2017, https://github.com/tobiasgf/lulu) to collapse aberrant and presumably erroneous MOTUs. The centroid sequences of each MOTU were compared using *ecotag* to obtain taxonomic assignment against a custom reference database built with all sequences from Arachnida and Hexapoda from the BOLD database (Ratnasingham and Hebert, 2007; downloaded in June 2018) and the invertebrate and fungi files from release 133 of the nucleotide sequence database from the EMBL repository (Kulikova *et al.*,, 2004). At this point, 16S–‘long’ and 16S–‘short’ output files were merged. The script *owi_add_taxonomy* was used to complete the taxonomic information (from kingdom to order level) of the annotated MOTUs. In a parallel MiSeq run, a library was sequenced that was prepared with only one of the samples of this study and four controlled tag combinations. The proportion of reads that didn’t match any of the four combinations was 0.4 %. Therefore, the final dataset was denoised by deleting from each sample those MOTUs with a relative abundance of reads lower than 0.4 % of the total reads of the sample, and by deleting those with equal or less than 10 reads in total. The detailed bioinformatics pipeline with all programs and scripts used can be found in Supplementary Material.

### Statistical analysis

All analyses of community composition were conducted using the package ‘vegan’ v2.4.4 (Oksanen *et al.*, 2013) in R v3.3.3 (R Development Core Team, 2017). Unless stated otherwise, only MOTUs assigned to Arthropoda were considered. Visualization of the recovered community structure from the different Malaise trap sample types (ethanol or homogenized tissue) was based on a non-metric multidimensional scaling (NMDS) analysis of a Jaccard or a Bray-Curtis dissimilarity matrix. As the results were similar, we only show the results based on the Jaccard index here. To test for differences between the communities recovered from ethanol and tissue with both 16S and COI, we used a Permutation Analysis of Variance (PERMANOVA) with the function ‘adonis’ for analysis of dissimilarity. Rarefaction curves were obtained with the function ‘rarecurve’.

To examine how the morphological traits of individual taxa affected their detectability in the preservative ethanol versus insect homogenate, we scored two sets of characters for each arthropod family: size (large, medium, small) and degree of sclerotization of the cuticle (strong, medium, weak). We then asked two questions: First, how do the traits of the taxon (size, sclerotization) affect its *probability of detection* (presence/absence) in the two types of samples? Second, how do the traits of the taxon affect its *relative read count*.

To answer the first question, we used a generalized linear mixed effects model of presence as a function of fixed effects Sample Type (ethanol or tissue), Size and Sclerotization, along with all two and three-way interactions. To account for the paired design (where both types of samples were derived from the same Malaise trap catches), we defined Sample ID as a random effect. Since the response was binomial, we assumed a logit-link and binomially distributed error.

To answer the second question, we focused on the log-ratio between the read counts of the same taxon from the same sample, as recorded from ethanol versus tissue homogenate. In other words, we used *R=log{(n*_*E*_+*1)/(n*_*T*_+*1)}* as our response, where n_E_ is the taxon- and sample-specific read count from the preservative ethanol, and n_T_ is the taxon- and sample-specific read count from the tissue homogenate of the same sample. We then fit a generalized linear mixed-effects model of R as a function of Size, Sclerotization and their interaction, assuming an identity link and normally distributed errors. Following the principle of model reduction, we dropped the non-significant interaction term before fitting the final model (Size: F_2,74_=7.54, P=0.001; Sclerotization: F_2,74_=6.68, P=0.002; interaction Size×Sclerotization: F_4,70_=0.22, P=0.93). To visualize the estimated effects, we derived least-squares means from the fitted model. All models were fitted with SAS for Windows version 9.4 (SAS Institute Inc., Cary, NC, USA), procedure GLIMMIX, treating Size and Sclerotization as categorical class-level variables.

## Results

### Summary of raw reads and MOTUs results

The MiSeq run produced a total of 17.4M reads plus approximately 1.97M control reads of PhiX, of which 13.8M passed the quality filtering and could be assigned to a sample. The average number of reads per sample was 1.73M ± 732,702 (s.d.), with ethanol-16S yielding the most abundant reads (2.64M) and leaf litter-COI the least abundant reads (797,856). Clustering with SWARM produced 60,460 and 34,296 MOTUs for 16S and COI respectively, but after curation with LULU and denoising, this was reduced to 481 and 522 MOTUs, respectively, in the final dataset. From these, 430 and 432 MOTUs, respectively, could be assigned to Arthropoda.

The proportion of Arthropoda/non-Arthropoda reads was highly variable depending on the sample and genetic marker (Figure S2). For 16S, the average proportion of Arthropoda reads was 92.1 %, ranging from 59.7 % to 100 % and with no clear pattern among different types of samples. For COI, the value ranged from 0 % to 100 %, with the leaf litter samples almost entirely comprised by non-Arthropoda reads. Only for three humus samples did we detect any substantial proportion of arthropod reads (20 %), as compared to an average of 71.5 ± 12.0 % and 97.5 ± 7.5 % arthropod reads in ethanol and tissue samples, respectively. In all but one sample (16S-H2T2: humus sample from site 2, second time point, amplified with 16S) the rarefaction curves reached the stationary phase well before reaching the maximum number of reads (Figures S3 and S4), suggesting that sequencing depth was enough for recovering all unique PCR products.

### Comparison between markers

The different taxonomic depth at which the Arthropoda MOTUs were identified reflects differences in the completeness of the reference databases used (Figure 2). The 16S reference database consisted of only 63,325 unique sequences, while the COI database was an order of magnitude larger, consisting of 666,795 unique sequences. As a consequence, the number of MOTUs identified to lower taxonomic levels (species, genus and family) was considerably higher in the COI dataset (411 out of 432) than in the 16S dataset (315 out of 430).

**Figure 2.**
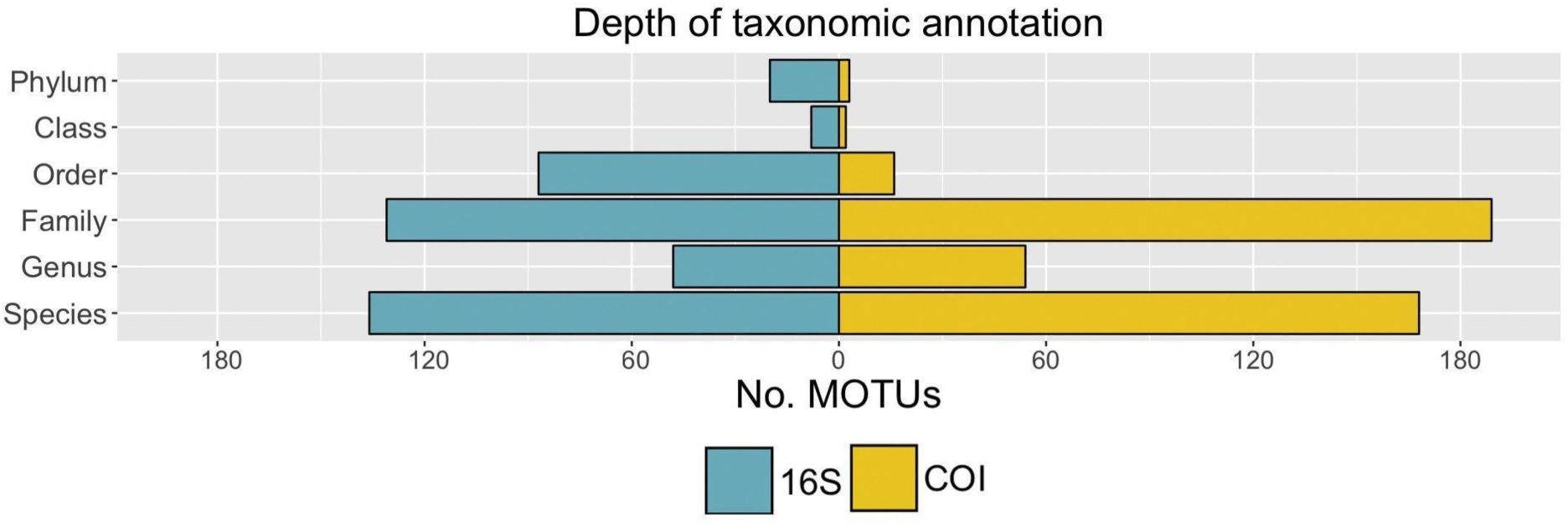
Number of MOTUs identified at each taxonomic level for 16S and COI when only Arthropoda OTUs are considered. The total number of arthropod MOTUs for 16S is 430 while for COI is 432.

For most taxa, COI recovered more MOTUs than 16S (Table S2). The number of MOTUs was equal for Hemiptera, Neuroptera, Ephemeroptera and Opiliones, while 16S recovered more MOTUs than COI for Diptera and Psocoptera. Representatives of Trichoptera, Symphypleona, Araneae, Mesostigmata and Sarcoptiformes were only recovered by COI, while MOTUs belonging to Orthoptera (family Acrididae) and Blattodea were only recovered by 16S.

The soil samples present a very different picture (Table S3). Most COI sequences belong to Fungi and other phyla and kingdoms (the unidentified and the non-Arthropoda sequences sum up to 96.37 % of the COI reads). Nevertheless, representatives of Orthoptera (family Tettigoniidae) and Araneae were only recovered by this marker. The rest of the orders except Symphypleona, Trombidiformes and Julida were exclusively or predominantly recovered by 16S.

At both species and genus level, most taxa were recovered exclusively by COI, with a lower proportion exclusively recovered by 16S, and even less taxa recovered by both markers simultaneously (Table 1). At the family and order levels, the majority of taxa were recovered by both markers, with a lower proportion detected exclusively by COI or 16S. These patterns are largely due to the (in)completeness of the reference libraries. When those taxa recovered by one marker but not present in the reference database of the other marker are eliminated, the situation changes: most species are recovered exclusively by 16S, and the overlap is higher for the genera, while the numbers for families and orders barely change.

**Table 1.** Taxa recovered exclusively by 16S or COI markers from the trap samples (*i.e.* bulk tissue or preservative fluid), or detected by both markers. The correction refers to eliminating from the reference database of one marker those taxa that were not present in the database of the other marker, with the intent of making detection failures comparable. From this table, we have explicitly excluded the soil samples, which were completely dominated by non-target DNA amplified by COI (see Figure S2).

| Level | Without Correction |  |  | With Correction |  |  |
| --- | --- | --- | --- | --- | --- | --- |
|  | 16S | COI | Common | 16S | COI | Common |
| **Species** | 84 | 136 | 29 | 59 | 14 | 29 |
| **Genus** | 53 | 131 | 49 | 53 | 33 | 49 |
| **Family** | 10 | 35 | 54 | 10 | 23 | 54 |
| **Order** | 3 | 5 | 11 | 3 | 5 | 11 |

In relation to the individual trap samples, at family level the overlap between 16S and COI is very close to 50 % – varying from 40 % to 64 % depending on the sample – (Table S4). Families recovered exclusively by COI represented around 20 % to 40 % of the detected families (with some exceptions). That is, depending on the sample, between 10 % to 30 % of the total family diversity detected for the target community was not recovered by COI.

### Trap samples versus soil samples

For both 16S and COI, the homogenized tissue samples were composed almost entirely of reads corresponding to Arthopoda (with exception of COI-T1T4, with approx. 75 % Arthropoda reads). While for 16S the ethanol samples showed a slight decrease in the proportion of reads corresponding to Arthropoda (with the exception of 16S-E2T2, which showed a larger decrease to approx. 60 %), for COI the proportion fell to 50-80 %. In the soil samples, the differences between COI and 16S are even more striking: for 16S, leaf litter and humus samples did not differ significantly from the ethanol in proportion of arthropod reads, while COI leaf litter samples were completely dominated by non-Arthropoda reads. The same applies to COI humus samples, with the exception of COI-H2T3, COI-H3T3 and COI-H3T4, which contained 25-30 % Arthropoda reads.

With respect to the number of MOTUs, again COI and 16S present more similar results for the trap samples than for the soil samples. The total number of MOTUs detected in the trap catches (Figure S5) was comparable between COI and 16S. The numbers decreased over time, presumably reflecting the temporal change in the insect communities. The MOTUs detected in preservative ethanol with COI were clearly fewer than those detected with 16S, due to a lower proportion of reads from arthropods. In the eDNA samples (both humus and leaf litter samples; Figure S6), 16S detected varying levels of MOTU richness depending on the sample, ranging from 9 to 31 MOTUs. COI only detected a small fraction of the arthropod diversity present in the samples; instead, the COI results were dominated by non-target DNA sequences (Table S3).

From the 430 MOTUs detected by 16S, 337 were found in the trap samples (summing ethanol and tissue) and 120 were detected in the soil (humus and leaf litter). Only 27 of these 430 MOTUs were detected both in the traps and the soil, with Diptera being the order with most MOTUs present in both sample types (18), followed by Coleoptera (3), and other orders with only one MOTU (Hemiptera, Psocoptera, Entomobryomorpha, unidentified Collembola and two unidentified Arthropoda). On each individual site, the overlap between soil and traps consisted of only a few MOTUs (Figure S7). COI detected only 14 MOTUs in the soil samples, out of the 432 found in total, and only two (one Coleoptera and one Symphypleona) appeared in both traps and soil. For COI, the total cumulative number of MOTUs for each site across the four time points (Figure 3) run very close to the curve of the traps, while with 16S, the total cumulative curve is separated from the curve of the traps, and reflects both the increase in both sample types. The increase of the total curve is the result of the sum of the trap and soil curves, indicating no transfer of MOTUs with time from the soil to the traps, which would be the case if the emergence of flying species from their ground-dwelling larval stages resulted in soil taxa appearing in the Malaise traps after some time lag.

**Figure 3.**
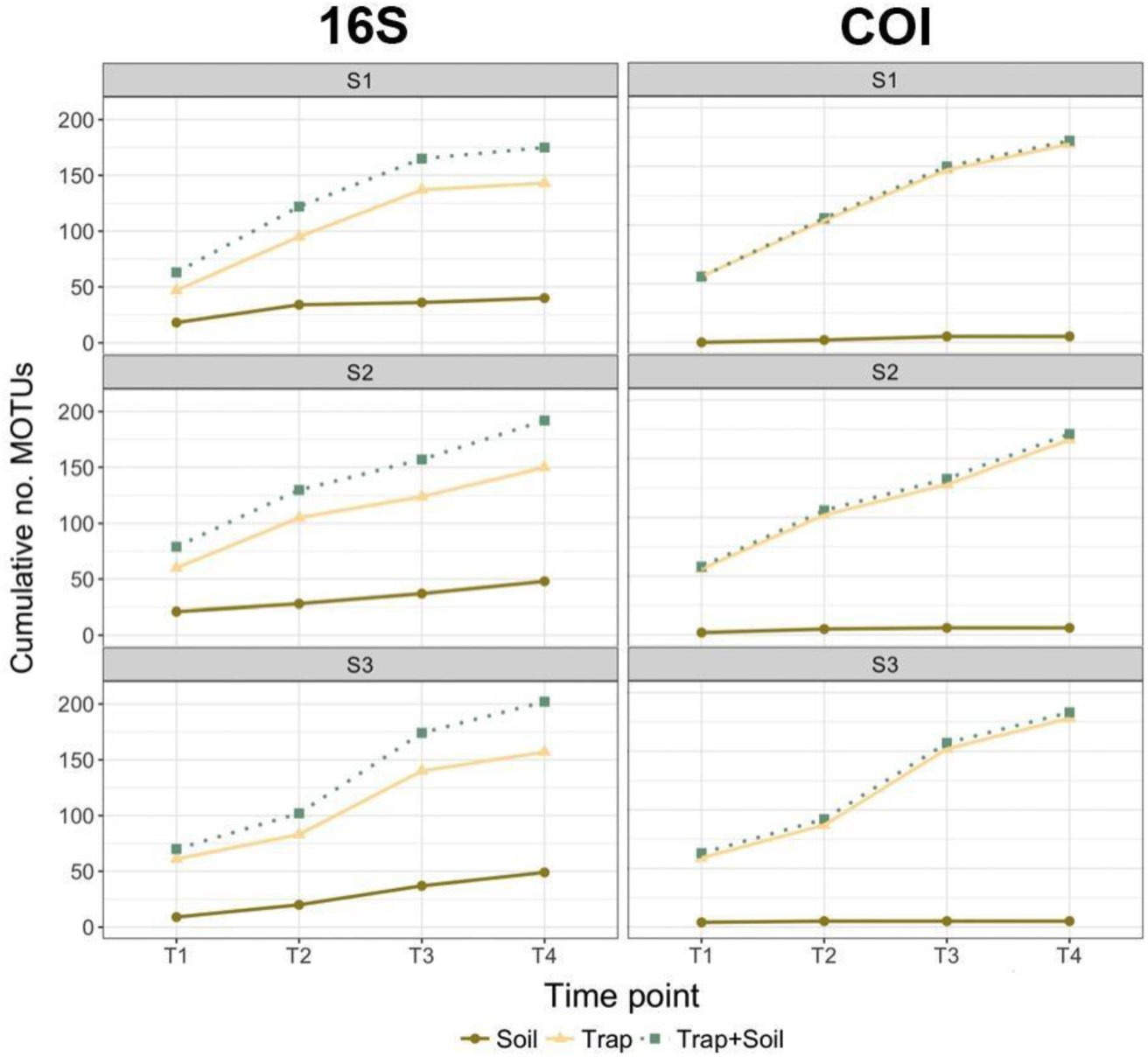
MOTU accumulation curves across the four sampling times for each trap, soil and combined trap + soil samples. Individual panels correspond to each site.

### Preservative ethanol versus homogenized trap samples

The community recovered from the Malaise trap samples was highly influenced by sample type (ethanol versus homogenate) independent of the marker used (ADONIS 16S ethanol–tissue: R^2^ = 0.12, P = 0.001; COI ethanol–tissue: R^2^ = 0.08, P = 0.001) (Figure 4). A closer look revealed very little overlap between the MOTUs recovered from tissue and ethanol in each sample, and in one case – for primer 16S S3T4 – no overlap at all (Figure S5). For 16S, the number of MOTUs recovered from ethanol was slightly higher than the number of MOTUs recovered from tissue (with some exceptions, *e.g.* S3T1), while for COI we observed the opposite pattern. This difference may have been caused by the higher proportion of non-Arthropoda sequences of COI detected in the ethanol samples (Figure S2).

**Figure 4.**
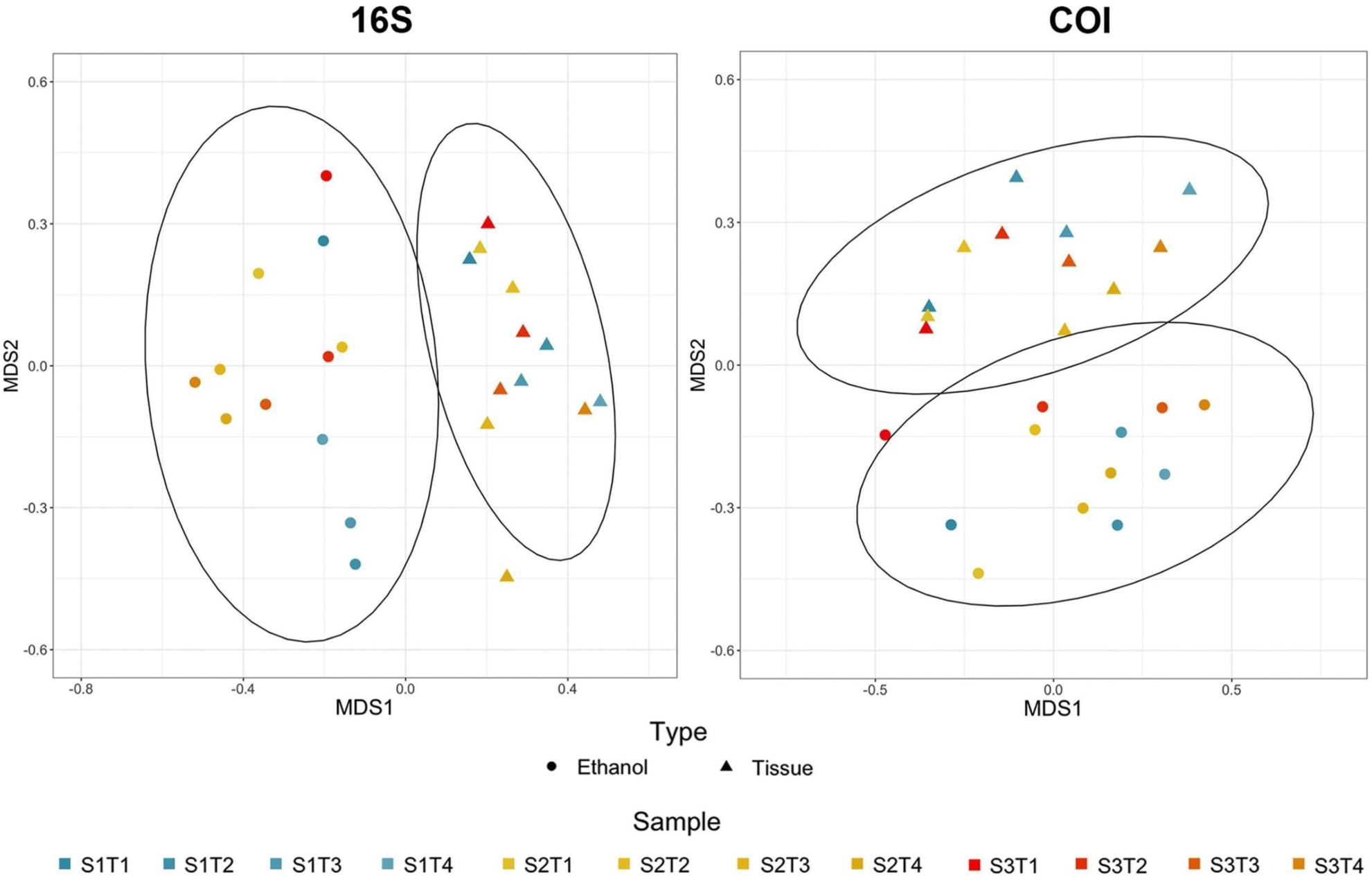
NMDS analysis of the recovered communities from ethanol and tissue for both 16S and COI markers, based on a Jaccard dissimilarity matrix.

Differences in community composition between the two sample types (homogenized tissue versus preservative ethanol) were correlated with insect traits. The probability with which a particular family was detected in a particular sample was significantly affected by size, sample type and sclerotization, with significant two-way interactions between all three factors (Table 2, with family-level patterns shown in Table S5). The effect of size was directly opposite for samples of different type: for ethanol samples, detection probability was highest for insects of small size, whereas for tissue homogenate, detection probability was lowest for small insects and more or less similar for insects of medium and large size (Figure 5, left). More highly sclerotized insects were less likely to be detected – but these differences were much smaller for samples of tissue homogenate than of preservative fluid (Figure 5, centre). Families of medium sclerotization and medium to large size were the most easily detected, with more complex patterns among other categories (Figure 5, right).

**Table 2.** The effect of Sample Type, Size and Sclerotization on the probability of detecting an insect family. Shown are Type III tests of fixed effects of the generalized linear mixed-effects model described in the main text.

| Effect | *F*-ratio | Num DF | Den DF | *P* |
| --- | --- | --- | --- | --- |
| **Sample Type** | 12.33 | 1 | 425 | 0.0005 |
| **Size** | 0.18 | 2 | 425 | 0.8328 |
| **Sclerotization** | 0.66 | 2 | 425 | 0.5175 |
| **Sclerotization\*Sample Type** | 11.86 | 2 | 425 | <0.0001 |
| **Size\*Sample Type** | 12.17 | 2 | 425 | <0.0001 |
| **Size\*Sclerotization** | 2.77 | 4 | 425 | 0.0272 |

**Figure 5.**
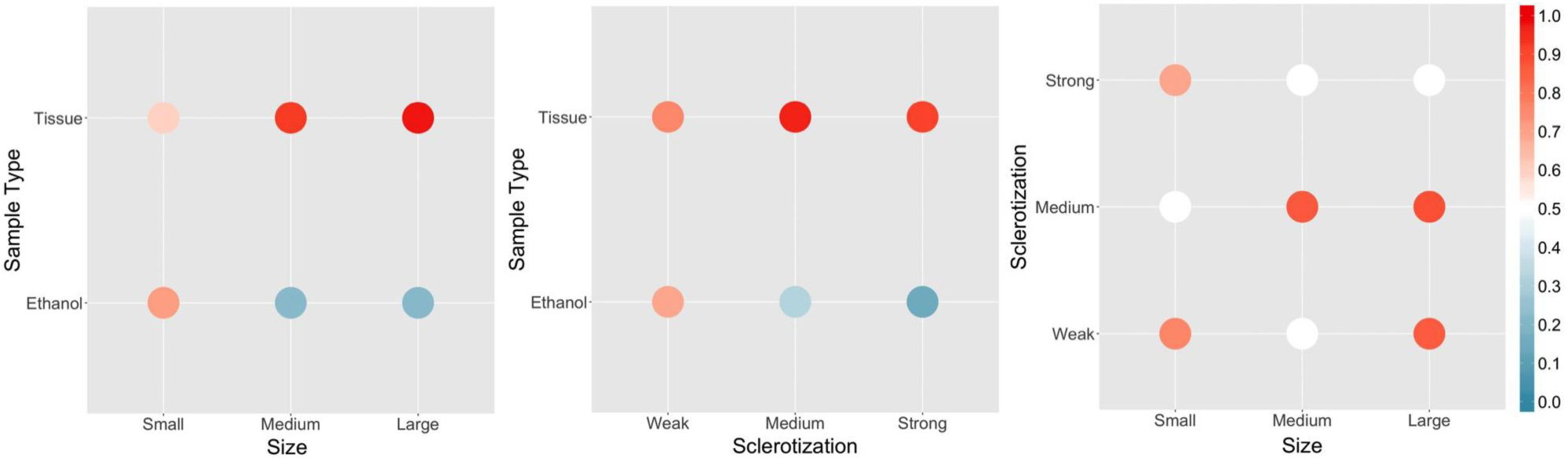
Effects of Sample Type, Size and Sclerotization on the probability of detecting an insect family, as derived from the generalized linear mixed-effects model summarized in Table 2.

Likewise, the relative read numbers recovered from samples of different types were strongly affected by insect traits, with both size and sclerotization having a significant effect. As there was no detectable interaction between the two factors (interaction Size×Sclerotization F_4,70_=0.22, P=0.93), this term was dropped from the final model, in which both Size (F_2,74_=7.54, P=0.001) and Sclerotization (F_2,74_=6.68, P=0.002) retained independent, significant effects. In brief, the log-ratio of counts from ethanol versus tissue declined with increasing size and sclerotization (Figure 6). In other words, the smaller and softer the insect family, the more abundant the reads from ethanol were relative to the reads from tissue.

**Figure 6.**
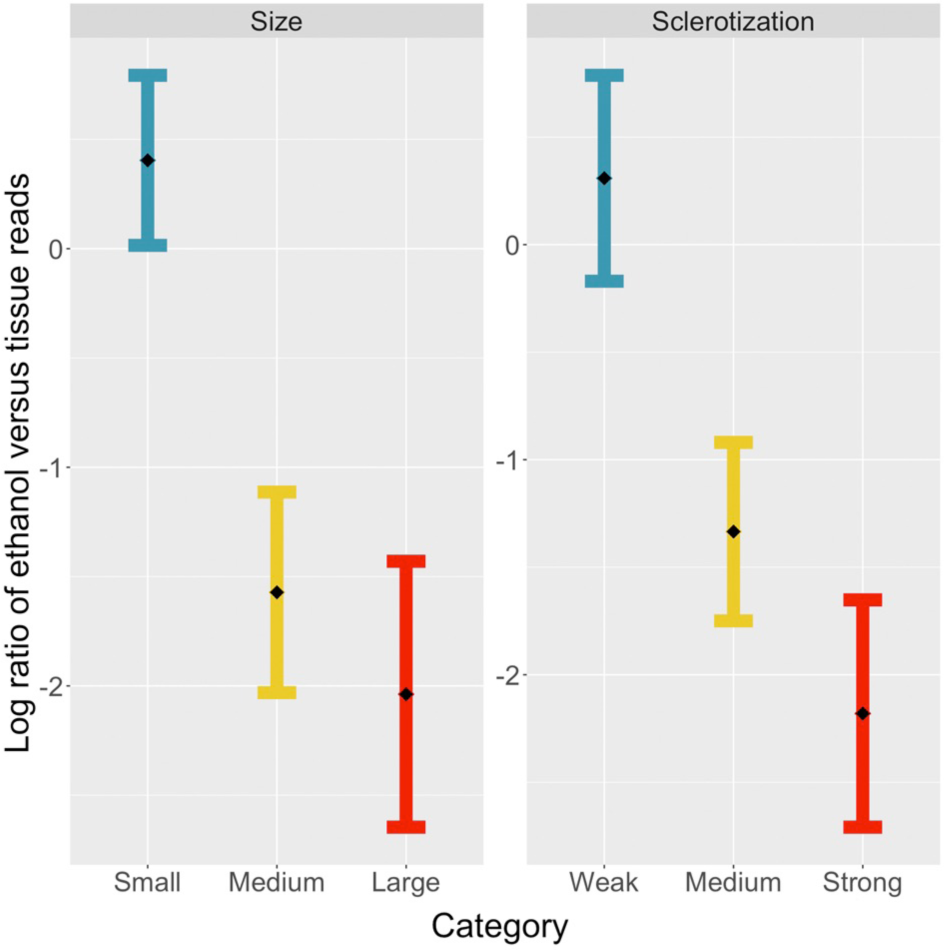
Least-squares means (with standard errors) from the fitted model of log-ratio between the read counts of the same taxon from the same sample, as recorded from ethanol versus tissue homogenate, as a function of the (left) size and (right) sclerotization of insect families.

## Discussion

In this study, we have surveyed total insect diversity at a selected site, comparing a widely used passive sampling device (a Malaise trap), with sampling of leaf litter and soil in the surroundings, as proposed by Yang *et al.*, (2014). In doing so, we also tested different approaches of suggested key impact on the metabarcoding process. On one hand, we compared the effects of the substrate analysed: an insect sample, or eDNA present in the soil. On the other hand, we compared analysis of DNA present in the insect tissue of the trap sample versus the DNA that had leaked out into the storage medium (ethanol) of the sample. We also explored analyses conducted with two primers optimized for insect metabarcoding: COI (a marker characterized by high variability and high capacity of species discrimination), and 16S (a less variable marker with broader taxonomic coverage but somewhat lower taxonomic resolution power), and their combination.

Overall, we find considerable differences between the results obtained with different markers and with different sampling protocols. The most striking result is that analysis of preservative ethanol appears to preferentially pick out small and soft-bodied insects that presumably comprise a small fraction of the total body mass of the trap sample, but that may comprise a significant fraction of the total specimen diversity. Our results show that the combined use of 16S and COI, as well as the metabarcoding of DNA from both the preservative ethanol and homogenized tissue, improves the recovery of arthropod diversity from Malaise trap catches. We also find that the highly degenerate primers required for taxonomically broad COI metabarcoding of insects are unsuitable for eDNA studies, simply because they pick up so many non-arthropod sequences. The alternative marker 16S recovered more arthropods in soil, but provided little overlap between both bulk and eDNA sample types. Below, we discuss these results in more detail.

### Comparison between markers

In terms of number of arthropod MOTUs, both markers were consistent, recovering 430 (16S) and 432 (COI) arthropods for the total dataset, respectively. The evenness in the number of arthropod MOTUs between both markers allows for an easy comparison of the quality of the taxonomic annotation allowed by each marker. As expected, MOTUs could be identified to a lower taxonomic level for COI than for 16S, although the difference was not as large as the difference in completeness of the reference databases. For COI, 95 % of MOTUs could be identified to family, genus or species, while this was true for only 73 % of 16S sequences. These results demonstrate that 16S has adequate taxonomic resolution for metabarcoding of insects, but they also confirms the necessity of creating a local reference database to obtain a biodiversity assessment of equal quality to that obtained by COI with this marker (Clarke *et al.*, 2014, Deagle *et al.*, 2014, Elbrecht *et al.*, 2016). However, the lack of reference sequences for 16S was not as problematic as the small size of the 16S reference library would have suggested. This may be because the Swedish insect fauna is fairly well known and has been subject to more 16S barcoding than the faunas of poorly documented, megadiverse regions of the world, such as the tropics. Possible drawbacks of 16S include the creation of spurious MOTUs with rRNA genes (or ‘taxonomic inflation’) caused by the difficulty of dealing with gaps in the bioinformatic processes (Clare *et al*., 2016), and increased chimera formation caused by the higher similarity between the sequences (Wilson *et al*., 2018). However, after dataset curation removing low abundance MOTUs, we obtained very similar numbers for COI and 16S, indicating that these problems can be circumvented by appropriate data cleaning approaches.

Even though 16S recovers fewer MOTUs than COI in most insect orders, 16S marker complements the results of COI, as has been concluded by several investigators (Deagle *et al.*, 2014, Clarke *et al.*, 2014, Marquina *et al.*, 2018), falsifying the hypothesis that a highly degenerate COI primer pair is enough to detect all biodiversity (Elbrecht *et al.*, 2016). To judge the overlap between the results obtained by the COI and 16S markers, it is necessary to focus on the taxa for which there is sufficient reference material to avoid biases caused by the vastly different sizes of the reference libraries. When this is done, the overlap is actually quite high (62 % at family level), and it is the 16S marker that recovers more exclusive taxa (59 species, 53 genera and 10 families) and not COI (14 species, 23 genera and 23 families). These numbers should nevertheless be treated with caution, as the incompleteness of the 16S database could have caused unrealistically low values for COI. The COI marker, even one associated with a highly degenerate primer pair like the one we used, is not sufficient to detect all insect biodiversity. Thus, our results are consistent with those of other studies advocating for complementary multi-marker approaches in metabarcoding of insect communities (Deagle *et al.*, 2014, Clarke *et al.*, 2014, Kaunisto *et al.*, 2017) and other taxa (Wangensteen *et al.*, 2018, Holman *et al.*, 2018).

### Preservative ethanol vs homogenized tissue from Malaise trap samples

Metabarcoding of homogenized insect tissue from trap samples comes at the cost of losing all the morphological information of the collected specimens. This prevents further study of the material, such as description of any potential new species from the material, or reference barcoding or detailed investigations of individual specimens. Therefore, analysis of preservative ethanol would open up many possibilities if it were an adequate replacement for analyses of tissue homogenate. Shokralla, Singer, & Hajibabaei (2010) were the first ones to achieve PCR amplification of DNA extracted directly from fixative ethanol from single specimen samples. Later, Hajibabaei *et al.*, (2012) did the same with freshwater benthic bulk samples, reporting high recovery success: 89 % of the taxa identified with the tissue extraction were also identified in the ethanol. Since then, studies testing the suitability of DNA metabarcoding of preservative ethanol have been scarce, which is surprising given the significant advantages of this method: in addition to the availability of the material for further sequencing or for taxonomic study, the method also comes with a much lower processing time in the laboratory than other non-destructive protocols, which tend to involve incubation of the sample in a lysis buffer. To our knowledge, there are only three studies of the performance of high-throughput sequencing from preservative ethanol of bulk samples subsequent to the two pioneering papers by Hajibabaei’s group. The first focused on genome skimming of mock samples of terrestrial and aquatic Coleoptera (Linard, Arribas, Andújar, Crampton-Platt, & Vogler 2017), while the second and third used metabarcoding of both mock and real samples from freshwater benthic macroinvertebrates (Erdozain *et al*., 2019; Zizka *et al.*, 2018). Our study is the first one to present data on DNA extracted from preservative ethanol from Malaise trap samples.

The methodology used to obtain the DNA from the ethanol varies between each of the previous studies: from simple evaporation of 1 ml of ethanol at 56 °C and resuspension in water (Erdozain *et al*., 2019; Hajibabaei *et al.*, 2012), through the more elaborate removal of the ethanol, including centrifugation to precipitate the DNA, drying of the precipitate and extraction with a commercial kit (Linard *et al.*, 2017), to passing through a filtering membrane which is incubated with a lysis buffer, in a similar way to what is done with the water in eDNA protocols (Zizka *et al.*, 2018). We chose the latter option to obtain all the free DNA and cells in the ethanol, rather than small aliquots that might have too low DNA concentration and from which some DNA sequences could be absent. Whether the choice of these three methodologies affects the detection of different taxa is unknown, and should be tested in the future.

We found that the insect community recovered from metabarcoding of preservative ethanol is strikingly different from the community recovered from homogenized tissue. Of the taxa we found, 36 families were exclusively found in ethanol, while 23 families were exclusively found in homogenates, and 17 were found in both substrates. The larger and more heavily sclerotized insects were recovered more frequently in the tissue samples, and the smaller and more weakly scleortized were primarly recovered in the ethanol. Nevertheless, some exceptions to this pattern can be pointed out. For instance, ladybugs (Coccinelidae, Coleoptera) are strongly sclerotized but are nevertheless found exclusively in preservative ethanol in some samples. This could be because they, when stressed, produce defensive excretions of which haemolymph is a main component, containing large amounts of DNA. Tipulids (Tipulidae, Diptera) are recovered by both the ethanol and the tissue, but their read numbers is almost three times larger in the ethanol. Although being large insects, tipulids have a very high surface to volume ratio, which could facilitate the entry of ethanol into the body and the leakage of DNA into the solvent. Another example is the oribatid mites (Oribatida, Sarcoptiformes), exclusively found in the ethanol samples. They have a strong tegument, but they can be so small that they were possibly not retained in the 0.6 mm pore size sieve, and therefore ended up in the ethanol filter rather than the tissue fraction of the sample. Also, some insects are known to vomit or regurgitate the intestinal contents when introduced alive in ethanol, and that can be the explanation of the exclusive presence of some families in preservative ethanol. However, without knowledge of the true taxonomic composition of the samples, it is impossible to say whether the results we observed are caused by any of these taxon-specific features or simply by PCR biases or other biases in DNA recovery during the processing of the sample. The absence of certain families of small insects in the tissue samples can be due to the fact that they represent only a small fraction of the total biomass, and hence of the DNA of the sample, as suggested by Elbrecht *et al*. (2017). However, the level of sclerotization still appears to play an important role, since most families of small but strongly sclerotized insects are preferentially found in the tissue samples.

In summary, our results contrast strongly with the findings of Hajibabaei *et al.*, (2012), who reported almost 100 % recovery in ethanol of the taxa found in tissue homogenate, but are in line with the results of Linard *et al.*, (2017), who were only able to recover 15 of 40 morphospecies from preservative ethanol versus 37 from the tissue, those of Zizka *et al.*, (2018), who found that 39 and 85 MOTUs out of 356 were only found in ethanol or homogenate, respectively, and the recent results from Erdozain *et al*. (2019), who found that metabarcoding of preservative ethanol consistently detected fewer taxa than morphological identification. As in the previous cases, detection probabilities were in general affected by the morphology of the insects. This could explain the positive results in Hajibabaei *et al.*, (2012), since their samples mostly consisted of larvae, which are weakly sclerotized in general and therefore should be well represented in preservative ethanol. In light of our results, it is clear that preservative ethanol-based DNA cannot be used as a non-destructive replacement of destructive tissue homogenization protocols in metabarcoding of Malaise trap samples. Instead, the two methods complement each other: preservative ethanol appears to retrieve small and weakly sclerotized insects, while homogenized tissue seems to be dominated by insect species with large total body mass.

### Malaise trap samples versus soil samples

Our results clearly show that COI is problematic in metabarcoding of environmental DNA samples like soil samples. This is mainly because it is amplified using highly degenerate primers, whose target/off-target amplification rate ratio is greatly affected by the proportion of DNA from non-target taxa in the sample. This effect explains the abundance of non-arthropod sequences in COI analyses of soil samples. It may also explain why COI analyses of preservative ethanol tend to pick up more non-target taxa than 16S analyses. To some extent, ethanol samples can be considered eDNA, since not only the DNA of the collected insects leaks out into the ethanol, but also the DNA of microorganisms (pathogens or symbionts) associated with the insects, and of those organisms that had an interaction with the collected specimen. A potential explanation for the predominance of fungal over insect sequences in the COI analyses of the soil samples could be that the fungi present in the soil are present alive while the environmental insect DNA is degraded. To deal with degradation, primers for shorter COI fragments have been designed with or without special focus on degraded eDNA (*e.g.* Vamos, Elbrecth, & Leese 2017; Zeale, Butin, Barker, Lees, & Jones 2011). Nevertheless, this explanation can probably be discarded due to the fact that the 16S amplicon, which is longer than the COI amplicon, was not affected. Thus, it can be concluded that the COI amplification of off-target sequences is caused by too high degeneracy of the primers and not by fragmentation of insect DNA.

For surveys focusing on total biodiversity in eDNA samples, highly degenerate primers can provide information from a very wide spectrum of taxa (Deiner *et al.*, 2016, Macher *et al.*, 2017, Majaneva *et al.*, 2018b). However, this comes at the expense of lower taxonomic coverage for any specific taxon, which may result in large parts of the community missing from the resulting reads. Therefore, for eDNA samples, it is often better to use less degenerate primers for more conserved markers — such as rRNA genes (Horton *et al.*, 2017) — even when this comes at the cost of reduced species discrimination capacity (Tang *et al.*, 2012, Wangesteen *et al.*, 2018). Our results show that the 16S primers chosen work quite well for analysis of insect community composition in eDNA samples, since the primer-binding site is more conserved than for COI, but the lower degeneracy prevents off-target amplification. At the same time, the fairly long amplicon read (350 bp) gives good taxonomic resolution (Marquina *et al*., 2018), better than that obtained with other 16S primers tried to date (Alberdi *et al.*, 2018, Clarke *et al.*, 2014, Elbrecht *et al.*,, 2016).

We find the limited overlap in MOTUs between soil and trap samples surprising for two reasons: first, we expected to find some soil inhabitant species that had climbed up and into the Malaise traps, and second, we expected to see a time-lagged overlap caused by ground-dwelling larvae detected in the soil samples, which – once developed into adults – would find their way into the traps. There were only a few examples of the first case, involving springtails and beetles found both in the soil and the trap samples. There were more examples of the second case, but still only 18 MOTUs, all belonging to Diptera (some of these cases could actually be explained by dead adults being detected in the soil samples). A possible reason for the absence of a time-lagged overlap between soil and trap catches is that both types of samples were taken late in the season (first samples were collected in early July), when most insects had probably already emerged. However, one may clearly expect insect communities found in different biological substrates in the same habitat or location to differ significantly (see, *e.g.*, Koziol *et al*., 2018).

## Conclusions

In this paper, we show that a multi-marker approach using COI and 16S markers enhances the recovery of biodiversity from insect bulk samples and eDNA. Our 16S marker does not appear to suffer from significantly lower taxonomic resolution than COI, unlike the nuclear rRNA markers that have been tried previously as complements to COI. This gain in resolution presumably emanates from the fact that our 16S marker is longer. Additionally, we find that the lower degeneracy of the primers amplifying 16S allows for on-target sequencing of the desired taxon in environmental samples, in contrast to the highly degenerate COI primers, which show low specificity and recover many phylogenetically distant non-target organisms. Thus, COI primers are not suitable for biodiversity surveys based on analysis of eDNA when only a narrow taxonomic focus is desired. Moreover, our results show that metabarcoding of DNA extracted from the preservative ethanol from Malaise samples is not equivalent to metabarcoding of the homogenized insect tissue from these samples. Thus, the former cannot be used as a non-destructive replacement for the latter. Nevertheless, metabarcoding of preservative ethanol clearly provides additional and valuable information, especially for small and soft insects, which appear to be underrepresented in the tissue homogenate. Therefore, analysis of preservative ethanol may be used in combination with analysis of tissue homogenate in order to obtain a more complete picture of the insect community than would be possible with only one of the methods alone.

## Acknowledgements

We are very thankful to Mattias Forshage, who provided valuable taxonomic information, and to Estela Barroso, for helping in the deployment and operation of the Malaise traps. This project was funded by the European Union’s Horizon 2020 research and innovation programme under the Marie Skłodowska–Curie grant agreement no. 642241 (BIG4 project, https://big4-project.eu) and by a grant from the Swedish Research Council (No. 2014-05901) to FR. The authors would like to acknowledge support from Science for Life Laboratory, the National Genomics Infrastructure, NGI, and Uppmax (Swedish National Infrastructure for Computing) for providing assistance in massive parallel sequencing and computational infrastructure.

## Author contributions

DM, TR and FR conceived and designed the study. DM collected the samples, and DM and RES carried out the laboratory work. DM did the bioinformatics processing and DM and TR performed the statistical analyses. DM wrote the first draft of the manuscript and prepared the figures and tables. All authors reviewed and contributed to subsequent drafts of the manuscript.

## Data accessibility

### Supplementary material

Additional supporting information may be found online in the Supporting Information section at the end of the article.

## References

Alberdi A., Aizpurua O., Gilbert M. T. P., & Bohmann K. (2018). Scrutinizing key steps for reliable metabarcoding of environmental samples. Methods in Ecology and Evolution, 9(1), 734–147. doi: 10.1111/2041-210X.12849

Andrews S. (2010). FastQC: a quality control tool for high-throughput sequence data. Available online at: http://www.bioinformatics.babraham.ac.uk/projects/fastqc

Andújar, C., Arribas, P., Yu, D. W., Vogler, A. P., & Emerson, B. C. (2018). Why the COI barcode should be the community DNA metabarcode for the metazoa. Molecular Ecology, 27(20), 3968–3975. doi: 10.1111/mec.14844

Aylagas, E., Mendibil, I., Borja, Á., & Rodríguez-Ezpeleta, N. (2016). Marine sediment sample pre-processing for macroinvertebrates metabarcoding: mechanical enrichment and homogenization. Frontiers in Marine Science, 3, 203. doi: 10.3389/fmars.2016.00203

Binladen, J., Gilbert, M. T. P., Bollback, J. P., Panitz, F., Bendixen, C., Nielsen, R., & Willerslev, E. (2007). The use of coded PCR primers enables high-throughput sequencing of multiple homolog amplification products by 454 parallel sequencing. PLoS ONE, 2(2), e197. doi: 10.1371/journal.pone.0000197

Bista, I., Carvalho, G. R., Tang, M., Walsh, K., Zhou, X., Hajibabaei, M., … & Christmas, M. (2018). Performance of amplicon and shotgun sequencing for accurate biomass estimation in invertebrate community samples. Molecular Ecology Resources, 18(5), 1020–1034. doi: 10.1111/1755-0998.12888

Bohmann, K., Evans, A., Gilbert, M. T. P., Carvalho, G. R., Creer, S., Knapp, M., … & De Bruyn, M. (2014). Environmental DNA for wildlife biology and biodiversity monitoring. Trends in Ecology & Evolution, 29(6), 358–367. doi: 10.1016/j.tree.2014.04.003

Boyer F., Mercier C., Bonin A., Le Bras Y., Taberlet P., & Coissac E. (2016). obitools: a unix-inspired software package for DNA metabarcoding. Molecular Ecology Resources, 16(1), 176–182. doi: 10.1111/1755-0998.12428

Clare, E. L., Chain, F. J., Littlefair, J. E., & Cristescu, M. E. (2016). The effects of parameter choice on defining molecular operational taxonomic units and resulting ecological analyses of metabarcoding data. Genome, 59(11), 981–990. doi: 10.1139/gen-2015-0184

Clarke L. J., Soubrier J., Weyrich L. S., & Cooper A. (2014). Environmental metabarcodes for insects: in silico PCR reveals potential for taxonomic bias. Molecular Ecology Resources, 14(6), 1160–1170. doi: 10.1111/1755-0998.12265

Cowart D. A., Pinheiro M., Mouchel O., Maguer M., Grall J., Miné J., & Arnaud-Haond S. (2015). Metabarcoding Is Powerful yet Still Blind: A Comparative Analysis of Morphological and Molecular Surveys of Seagrass Communities. PLoS ONE, 10(2), e0117562. doi: 10.1371/journal.pone.0117562

Creer, S., Deiner, K., Frey, S., Porazinska, D., Taberlet, P., Thomas, W. K., … & Bik, H. M. (2016). The ecologist’s field guide to sequence□based identification of biodiversity. Methods in Ecology and Evolution, 7(9), 1008–1018. doi: 10.1111/2041-210X.12574

Deagle B. E., Jarman S. N., Coissac E., Pompanon F., & Taberlet P. (2014). DNA metabarcoding and the cytochrome c oxidase subunit I marker: not a perfect match. Biology Letters, 10(9), 20140562–20140562. doi: 10.1098/rsbl.2014.0562

Deiner K., Fronhofer E. A., Mächler E., Walser J.-C., & Altermatt, F. (2016). Environmental DNA reveals that rivers are conveyer belts of biodiversity information. Nature Communications, 7, 12544. doi: 10.1038/ncomms12544

Deiner, K., Bik, H. M., Mächler, E., Seymour, M., Lacoursière□Roussel, A., Altermatt, F., … & Pfrender, M. E. (2017). Environmental DNA metabarcoding: transforming how we survey animal and plant communities. Molecular Ecology, 26(21), 5872–5895. doi: 10.1111/mec.14350

Drummond, A. J., Newcomb, R. D., Buckley, T. R., Xie, D., Dopheide, A., Potter, B. C., … & Park, D. (2015). Evaluating a multigene environmental DNA approach for biodiversity assessment. GigaScience, 4(1), 46. doi: 10.1186/s13742-015-0086-1

Elbrecht V., & Leese F. (2017). Validation and development of COI metabarcoding primers for freshwater macroinvertebrate bioassessment. Frontiers in Environmental Science, 5, 11. doi: 10.3389/fenvs.2017.00011

Elbrecht V., Peinert B., & Leese F. (2017). Sorting things out: Assessing effects of unequal specimen biomass on DNA metabarcoding. Ecology and Evolution, 7(17), 6918–6926. doi: 10.1002/ece3.3192

Elbrecht V., Taberlet P., Dejean T., Valentini A., Usseglio-Polatera P., Beisel J.-N., Coissac E., Boyer F., & Leese F.(2016). Testing the potential of a ribosomal 16S marker for DNA metabarcoding of insects. PeerJ, 4(4), e1966–12. doi: 10.7717/peerj.1966

Emilson, C. E., Thompson, D. G., Venier, L. A., Porter, T. M., Swystun, T., Chartrand, D., … & Hajibabaei, M. (2017). DNA metabarcoding and morphological macroinvertebrate metrics reveal the same changes in boreal watersheds across an environmental gradient. Scientific Reports, 7(1), 12777. doi: 10.1038/s41598-017-13157-x

Epp, L. S., Boessenkool, S., Bellemain, E. P., Haile, J., Esposito, A., Riaz, T., … & Stenøien, H. K. (2012). New environmental metabarcodes for analysing soil DNA: potential for studying past and present ecosystems. Molecular Ecology, 21(8), 1821–1833. doi: 10.1111/j.1365-294X.2012.05537.x

Erdozain, M., Thompson, D. G., Porter, T. M., Kidd, K. A., Kreutzweiser, D. P., Sibley, P. K., … & Hajibabaei, M. (2019). Metabarcoding of storage ethanol vs. conventional morphometric identification in relation to the use of stream macroinvertebrates as ecological indicators in forest management. Ecological Indicators, 101, 173–184. doi: 10.1016/j.ecolind.2019.01.014

Freeland J. R. (2017). The importance of molecular markers and primer design when characterizing biodiversity from environmental DNA. Genome, 60(4), 358–374. doi: 10.1139/gen-2016-0100

Frøslev T. G., Kjøller R., Bruun H. H., Ejrnæs R., Brunbjerg A. K., Pietroni C., & Hansen A. J. (2017). Algorithm for post-clustering curation of DNA amplicon data yields reliable biodiversity estimates. Nature Communications, 8(1), 1188. doi: 10.1038/s41467-017-01312-x

Gibson J. F., Shokralla S., Porter T. M. King, I. van Konynenburg S., Janzen D. H., Hallwachs W., & Hajibabaei M. (2014). Simultaneous assessment of the macrobiome and microbiome in a bulk sample of tropical arthropods through DNA metasystematics. Proceedings of the National Academy of Sciences, 111(22), 8007–8012. doi: 10.1073/pnas.1406468111

Gibson, J. F., Shokralla, S., Curry, C., Baird, D. J., Monk, W. A., King, I., & Hajibabaei, M. (2015). Large-scale biomonitoring of remote and threatened ecosystems via high-throughput sequencing. PLoS ONE, 10(10), e0138432–15. doi: 10.1371/journal.pone.0138432

Hajibabaei M., Spall J. L., Shokralla S., & van Konynenburg S. (2012). Assessing biodiversity of a freshwater benthic macroinvertebrate community through non-destructive environmental barcoding of DNA from preservative ethanol. BMC Ecology, 12(1), 1–1. doi: 10.1186/1472-6785-12-28

Holman L. E., de Bruyn M., Creer S., Carvalho G., Robidart J., & Rius, M. (2018). The detection of novel and resident marine non-indigenous species using environmental DNA metabarcoding of seawater and sediment. bioRxiv, 440768. https://doi.org/10.1101/440768

Horton D. J., Kershner M. W., & Blackwood C. B. (2017). Suitability of PCR primers for characterizing invertebrate communities from soil and leaf litter targeting metazoan 18S ribosomal or cytochrome oxidase I (COI) genes. European Journal of Soil Biology, 80, 43–48. doi: 10.1016/j.ejsobi.2017.04.003

Kaunisto K. M., Roslin T., Sääksjärvi I. E., & Vesterinen E. J. (2017). Pellets of proof: First glimpse of the dietary composition of adult odonates as revealed by metabarcoding of feces. Ecology and Evolution, 7(20), 8588–8598. doi: 10.1002/ece3.3404

Kocher, A., Gantier, J. C., Gaborit, P., Zinger, L., Holota, H., Valiere, S., … & Murienne, J. (2017). Vector soup: high-throughput identification of Neotropical phlebotomine sand flies using metabarcoding. Molecular Ecology Resources, 17(2), 172–182. doi: 10.1111/1755-0998.12556

Koziol, A., Stat, M., Simpson, T., Jarman, S., DiBattista, J. D., Harvey, E. S., … & Bunce, M. (2018). Environmental DNA metabarcoding studies are critically affected by substrate selection. Molecular Ecology Resources. doi:https://doi.org/10.1111/1755-0998.12971

Krehenwinkel H., Wolf M., Lim J. Y., Rominger A. J., Simison W. B., & Gillespie R. G. (2017). Estimating and mitigating amplification bias in qualitative and quantitative arthropod metabarcoding. Scientific reports, 7, 17668. doi: 10.1038/s41598-017-17333-x

Krehenwinkel H., Fong M., Kennedy S., Huang E. G., Noriyuki S., Cayetano L., & Gillespie R. G. (2018). The effect of DNA degradation bias in passive sampling devices on metabarcoding studies of arthropod communities and their associated microbiota. PLoS ONE, 13(1), e0189188. doi: 10.1371/journal.pone.0189188

Kulikova, T., Aldebert, P., Althorpe, N., Baker, W., Bates, K., Browne, P., … & Faruque, N. (2004). The EMBL nucleotide sequence database. Nucleic Acids Research, 32(suppl_1), D27–D30. doi: 10.1093/nar/gki098

Linard B., Arribas P., Andújar C., Crampton-Platt A., & Vogler A. P. (2016). Lessons from genome skimming of arthropod-preserving ethanol. Molecular Ecology Resources, 16(6), 1365–1377. doi: 10.1111/1755-0998.12539

Macher J.-N., Vivancos A., Piggott J. J., Centeno F. C., Matthaei C. D., & Leese, F. (2018). Comparison of environmental DNA and bulk-sample metabarcoding using highly degenerate cytochrome coxidase I primers. Molecular Ecology Resources, 18(6), 1456–1468. doi: 10.1111/1755-0998.12940

Mahé F., Rognes T., Quince C., de Vargas C, & Dunthorn M. (2015). Swarm v2: highly-scalable and high-resolution amplicon clustering. PeerJ, 3, e1420 doi: 10.7717/peerj.1420

Majaneva, M., Diserud, O. H., Eagle, S. H. C., Hajibabaei, M., & Ekrem, T. (2018a). Choice of DNA extraction method affects DNA metabarcoding of unsorted invertebrate bulk samples. Metabarcoding and Metagenomics, 2, e26664. doi: 10.3897/mbmg.2.26664

Majaneva M., Diserud O. H., Eagle S. H. C., Boström E., Hajibabaei M., & Ekrem, T. (2018b). Environmental DNA filtration techniques affect recovered biodiversity. Scientific Reports, 8:4682. doi: 10.1038/s41598-018-23052-8

Marquina D., Andersson A. F., & Ronquist F. (2018). New mitochondrial primers for metabarcoding of insects, designed and evaluated using in silico methods. Molecular Ecology Resources, 19(1), 90–104. doi: 10.1111/1755-0998.12942

Morinière, J., de Araujo, B. C., Lam, A. W., Hausmann, A., Balke, M., Schmidt, S., … & Haszprunar, G. (2016). Species Identification in Malaise Trap Samples by DNA Barcoding Based on NGS Technologies and a Scoring Matrix. PLoS ONE, 11(5), e0155497. doi: 10.1371/journal.pone.0155497

Oksanen, J., Blanchet, F. G., Kindt, R., Legendre, P., Minchin, P. R., O’hara, R. B., … & Oksanen, M. J. (2013). Package Vegan: Community ecology package, version 2.0 10.

Porter, T. M., & Hajibabaei, M. (2018). Scaling up: A guide to high-throughput genomic approaches for biodiversity analysis. Molecular Ecology, 27(2), 313–338. doi: 10.1111/mec.14478

R Development Core Team (2017). R: a Language and Environment for Statistical Computing R Foundation for Statistical Computing. Austria.

Ratnasingham S., & Hebert P. D. (2007). BOLD: The Barcode of Life Data System (http://www.barcodinglife.org). Molecular Ecology Notes, 7(3), 355–364. doi: 10.1111/j.1471-8286.2007.01678.x

Rognes T., Flouri T., Nichols B., Quince C., & Mahé F. (2016). VSEARCH: a versatile open source tool for metagenomics. PeerJ, 4, e2584. doi: 10.7717/peerj.2584

Shokralla S., Porter T. M., Gibson J. F., Dobosz R., Janzen D. H., Hallwachs W., Golding G. B., & Hajibabaei M. (2015). Massively parallel multiplex DNA sequencing for specimen identification using an Illumina MiSeq platform. Scientific Reports, 5, 9687. doi: 10.1038/srep09687

Shokralla S., Singer G. A. C., & Hajibabaei M. (2010). Direct PCR amplification and sequencing of specimens’ DNA from preservative ethanol. BioTechniques, 48(3), 233–234. doi: 10.2144/000113362

Tang C. Q., Leasi F., Obertegger U., Kieneke A., Barraclough T. G., & Fontaneto D. (2012). The widely used small subunit 18S rDNA molecule greatly underestimates true diversity in biodiversity surveys of the meiofauna. Proceedings of the National Academy of Sciences, 109(40), 16208–162012. doi: 10.1073/pnas.1209160109/-/DCSupplemental

Vamos E., Elbrecht V., & Leese F. (2017). Short COI markers for freshwater macroinvertebrate metabarcoding. Metabarcoding and Metagenomics, 1(5), e14625–20. doi: 10.3897/mbmg.1.14625

Wangensteen O. S., Palacín C., Guardiola M., & Turon X. (2018). DNA metabarcoding of littoral hard-bottom communities: high diversity and database gaps revealed by two molecular markers. PeerJ, 6, e4705. doi: 10.7717/peerj.4705

Wilson, J. J., Brandon-Mong, G. J., Gan, H. M., & Sing, K. W. (2018). High-throughput terrestrial biodiversity assessments: mitochondrial metabarcoding, metagenomics or metatranscriptomics?. Mitochondrial DNA Part A. doi: 10.1080/24701394.2018.1455189

Yang C., Wang X., Miller J. A., de Blécourt M., Ji Y., Yang C., Harrison R. D., & Yu D. W. (2014). Using metabarcoding to ask if easily collected soil and leaf-litter samples can be used as a general biodiversity indicator. Ecological Indicators, 46, 379–389. doi: 10.1016/j.ecolind.2014.06.028

Yu D. W., Ji Y., Emerson B. C., Wang X., Ye C., Yang C., & Ding Z. (2012). Biodiversity soup: metabarcoding of arthropods for rapid biodiversity assessment and biomonitoring. Methods in Ecology and Evolution, 3(4), 613–623. doi: 10.1111/j.2041-210X.2012.00198.x

Zeale M. R., Butlin R. K., Barker G. L. A., Lees D. C., & Jones G. (2011) Taxon-specific PCR for DNA barcoding arthropod prey in bat faeces. Molecular Ecology Resources, 11(2), 236–244. doi: 10.1111/j.1755-0998.2010.02920.x

Zizka V. M., Leese F., Peinert B., & Geiger M. F. (2018). DNA metabarcoding from sample fixative as a quick and voucher preserving biodiversity assessment method. Genome.doi:https://doi.org/10.1139/gen-2018-0048

