## Supplementary material for "Establishing insect community composition using metabarcoding of soil samples, and preservative ethanol and homogenate from Malaise trap catches: surprising inconsistencies between methods": Supplemetal material

**Table S5.** List of families recovered exclusively from ethanol or tissue, or both from ethanol and tissue, ordered on degree of sclerotization and size (hard and large at the top, small and weak at the bottom). A blue square indicates when representatives of the family were detected in ethanol, and a red square when they were detected in tissue. The number of reads from each sample type is indicated in the rightmost two columns.

| | | | ETHANOL | | | | | | | | | | | | TISSUE | | | | | | | | | | | | | |
|---|---|---|---|---|---|---|---|---|---|---|---|---|---|---|---|---|---|---|---|---|---|---|---|---|---|---|---|---|
| | | | SITE 1 | | | | SITE 2 | | | | SITE 3 | | | | SITE 1 | | | | SITE 2 | | | | SITE 3 | | | | Number of reads | |
| **Family** | **Size** | **Sclerot.** | 1 | 2 | 3 | 4 | 1 | 2 | 3 | 4 | 1 | 2 | 3 | 4 | 1 | 2 | 3 | 4 | 1 | 2 | 3 | 4 | 1 | 2 | 3 | 4 | **Ethanol** | **Tissue** |
| Cerambycidae | Large | Strong | | | | | | | | | | | | | | | | | ■ | | | | ■ | | | | 0 | 393 |
| Latridiidae | Large | Strong | | | | | | | | | | | | | | | | ■ | | | | | | | | | 0 | 218 |
| Acanthosomatidae | Large | Strong | | | | | | | | | | | | | | ■ | ■ | ■ | | ■ | | ■ | | | | | 0 | 86 974 |
| Pentatomidae | Large | Strong | | | | | | | | | | | | | | | | | | | | | | ■ | | | 0 | 1 989 |
| Acrididae | Large | Strong | | | | | | | | | | ■ | | | | | | | | | ■ | | | | | | 517 | 1 997 |
| Attelabidae | Medium | Strong | | | | | | | | | | | | | | | | | | | ■ | | | | | | 0 | 5 888 |
| Curculionidae | Medium | Strong | | | | | | | | | | | | | ■ | | | | | | | | | | | | 0 | 118 |
| Elateridae | Medium | Strong | | | | | | | | | | | | | | | | | ■ | | | | ■ | | | | 0 | 6 835 |
| Leiodidae | Medium | Strong | | | | | | | | | | | | | | | | | | | | | | | | ■ | 0 | 176 |
| Melandryidae | Medium | Strong | | | | | | | | | | | | | | | | | | | | | ■ | ■ | | | 0 | 208 |
| Lygaeidae | Medium | Strong | | | | | | | | | | | | | | | | ■ | | | | | | | | | 0 | 1 553 |
| Ichneumonidae | Medium | Strong | | | | | | | | | | | | | ■ | ■ | ■ | ■ | ■ | ■ | ■ | ■ | ■ | ■ | ■ | ■ | 0 | 100 890 |
| Coccinellidae | Small | Strong | ■ | | ■ | | ■ | ■ | ■ | ■ | ■ | ■ | ■ | | | | ■ | | | | ■ | ■ | ■ | | ■ | | 51 574 | 29 554 |
| Cryptophagidae | Small | Strong | ■ | | | | | | | | | | | | ■ | | | | | | | ■ | | | | | 711 | 935 |
| Nitidulidae | Small | Strong | | | | | | | | | | | | | | | ■ | | | | | | | | | | 0 | 2 612 |
| Throscidae | Small | Strong | ■ | | | | | | | | | | | | ■ | ■ | ■ | | | ■ | ■ | | | | | | 3 006 | 8 330 |
| Scirtidae | Small | Strong | | | | | | | | | | | | | | | ■ | | | | | | | | | | 0 | 147 |
| Oribatulidae | Small | Strong | | | | | | | | | | | | ■ | | | | | | | | | | | | | 2 139 | 0 |
| Asilidae | Large | Medium | | | | | | | | | | ■ | | | | ■ | | | | ■ | | | | ■ | | | 932 | 4 293 |
| Calliphoridae | Large | Medium | | | | | | | | | | | | | | | | | | ■ | | ■ | ■ | ■ | ■ | ■ | 0 | 20 582 |
| Tabanidae | Large | Medium | | | | | | | | | | | | | | | | | ■ | | | | ■ | | | | 0 | 5 507 |
| Tachinidae | Large | Medium | | | | | | | | | | | | | | | | | ■ | ■ | | | | | | | 0 | 2 970 |
| Therevidae | Large | Medium | | | | | | | | | | | | | ■ | | | | | | | | | | | | 0 | 278 |
| Tipulidae | Large | Medium | ■ | | | | | ■ | ■ | | | ■ | | | ■ | | | | | ■ | ■ | | | | | ■ | 29 837 | 8 325 |
| Phalangiidae | Large | Medium | | | | | | | | | | | | | | | ■ | ■ | | | | | | | | | 0 | 6 963 |
| Ectobiidae | Medium | Medium | | | | | | | ■ | | ■ | | | | | | | | | | ■ | | | | | | 5 343 | 2 698 |
| Anisopodidae | Medium | Medium | | | | | | | | | | | | | | | | ■ | | | | | | | | | 0 | 2 587 |
| Anthomyiidae | Medium | Medium | | | | | | | | | | | | | ■ | ■ | ■ | | ■ | ■ | ■ | | ■ | ■ | ■ | ■ | 0 | 26 997 |
| Conopidae | Medium | Medium | | | | | | | | | | | | | | | | | | | | | | ■ | | | 0 | 530 |
| Dolichopodidae | Medium | Medium | ■ | ■ | | | ■ | | | | ■ | ■ | | | ■ | ■ | | | ■ | | | | ■ | ■ | | | 110 540 | 36 132 |
| Lauxaniidae | Medium | Medium | | | | | | | | | | | | | | | ■ | | | | | | | | | | 0 | 487 |
| Mycetophilidae | Medium | Medium | | | | ■ | | | | ■ | | | | | | | ■ | ■ | | | | ■ | | | | | 9 783 | 10 630 |
| Rhagionidae | Medium | Medium | | | | | | | | | | | | | | ■ | | | | ■ | | | | | | | 0 | 1 036 |
| Scathophagidae | Medium | Medium | | | | | | | | | | | | | | | | | | | ■ | | | | | | 0 | 837 |
| Sciomyzidae | Medium | Medium | | | | | | | | | | | | | ■ | | | | ■ | | | | | | | | 0 | 1 099 |
| Miridae | Medium | Medium | | ■ | | | ■ | | | | | | | | | | | | | | | | | | | | 6 714 | 0 |
| Braconidae | Medium | Medium | | ■ | | | | | | | | | | | | | | | | | | ■ | | | | ■ | 471 | 1 780 |
| Tenthredinidae | Medium | Medium | | | | | | | | | | | | | | ■ | | | | ■ | | | | | | | 0 | 700 |
| Araneidae | Small | Medium | | | | | | | ■ | | | | | | | | | | | | | | | | | | 3 569 | 0 |
| Linyphiidae | Small | Medium | | | | | | | | | | | | | | | | ■ | | | | ■ | | | | | 0 | 1 657 |
| Salticidae | Small | Medium | | | | | | | | | | ■ | | | | | | | | | | | | | | | 216 | 0 |
| Tetragnathidae | Small | Medium | | | | | | | | | | | | | | | | ■ | | | | ■ | | | | | 0 | 3 493 |
| Heleomyzidae | Small | Medium | | | | | | | | | | | | | | | ■ | | | | ■ | ■ | | | | | 0 | 4 577 |
| Hybotidae | Small | Medium | | ■ | | | | | | | | ■ | | | | | | | | | | | | | | | 3 509 | 0 |
| Melicharidae | Small | Medium | | | | | | | | | | | ■ | | | | | | | | | | | | | | 1 108 | 0 |
| Parasitidae | Small | Medium | | | | | | | | | | | | | | | | ■ | | | | | | | | | 0 | 261 |
| Phytoseiidae | Small | Medium | | | ■ | | | | | | | | | | | | | | | | | | | | | | 561 | 0 |
| Limnephilidae | Small | Medium | | | | | | | | | | | | | | ■ | | | | | | | | | | | 0 | 102 |
| Geometridae | Large | Weak | ■ | ■ | | | | ■ | | ■ | ■ | ■ | | ■ | ■ | ■ | | | ■ | ■ | ■ | ■ | ■ | ■ | ■ | ■ | 51 616 | 59 735 |
| Noctuidae | Large | Weak | | | | | | ■ | | | ■ | ■ | | | ■ | | | | ■ | ■ | | | ■ | ■ | ■ | | 2 450 | 69 289 |

| | | | | | | | | | | | | | | | | | | | | | | | | | | | | |
|---|---|---|---|---|---|---|---|---|---|---|---|---|---|---|---|---|---|---|---|---|---|---|---|---|---|---|---|---|
| Cantharidae | Medium | Weak | | | | | | | | | | | | | | ■ | | | ■ | | | | | | | | 0 | 1 964 |
| Keroplatidae | Medium | Weak | | | | | | ■ | | | | | | | | | | | | | | | | | | | 352 | 0 |
| Baetidae | Medium | Weak | | | | | | | | | | ■ | | | | | | | | | | | | | | | 4 659 | 0 |
| Crambidae | Medium | Weak | | | | | | | | | | | | | | ■ | ■ | | | | | | | | | | 0 | 1 562 |
| Psychidae | Medium | Weak | ■ | | | | | | | | | | | | | | | | | | | | | | | ■ | 276 | 195 |
| Cecidomyiidae | Small | Weak | ■ | ■ | ■ | ■ | ■ | ■ | ■ | ■ | ■ | ■ | ■ | ■ | ■ | ■ | ■ | ■ | | | | | | | ■ | ■ | 392 129 | 5 433 |
| Chironomidae | Small | Weak | ■ | ■ | ■ | | ■ | ■ | ■ | ■ | ■ | ■ | ■ | | | | ■ | | ■ | | ■ | | | | | | 38 131 | 11 900 |
| Psychodidae | Small | Weak | | | | ■ | ■ | ■ | | | | | | ■ | | | | | | | | | | | | | 1 941 | 0 |
| Sciaridae | Small | Weak | ■ | ■ | ■ | ■ | ■ | ■ | ■ | ■ | | ■ | ■ | ■ | | ■ | ■ | ■ | | ■ | ■ | ■ | | ■ | ■ | ■ | 95 671 | 14 052 |
| Entomobryidae | Small | Weak | ■ | | | ■ | ■ | ■ | ■ | ■ | | ■ | ■ | ■ | | | | | | | | | | | | | 231 599 | 0 |
| Aphididae | Small | Weak | ■ | ■ | | | | | | | | | | | ■ | | | | | | | | | | | | 4 799 | 348 |
| Lachnidae | Small | Weak | | | | | | | | | | | | | | | | | | | ■ | | | | | | 0 | 1 612 |
| Adelidae | Small | Weak | ■ | | | | | | | | | | | | | | | | | | | | | | | | 301 | 0 |
| Coleophoridae | Small | Weak | | ■ | | | | | | | | | | | | | | | | | | | | | | | 1 204 | 0 |
| Cosmopterigidae | Small | Weak | | | | | | | | | | ■ | | | | | | | | | | | | | | | 910 | 0 |
| Gracillariidae | Small | Weak | | | ■ | ■ | | | | | | ■ | | ■ | | | | | | | | | | | | | 823 | 0 |
| Incurvariidae | Small | Weak | ■ | | | | | | | | | | | | | | | | | | | | | | | | 1 462 | 0 |
| Oecophoridae | Small | Weak | | | | | ■ | | | | | | | | | | | | ■ | | | | | | | | 4 443 | 401 |
| Pyralidae | Small | Weak | | | | | | | | | | | | | | | | | | | | | | ■ | | | 0 | 84 |
| Yponomeutidae | Small | Weak | | | | | ■ | | | | | | | | | | | | | | | | | | | | 11 820 | 0 |
| Coniopterygidae | Small | Weak | | ■ | | | | ■ | | | | | | | | | | | | | | | | | | | 2 890 | 0 |
| Peripsocidae | Small | Weak | ■ | | ■ | ■ | ■ | | ■ | ■ | ■ | | ■ | | ■ | | ■ | | ■ | | ■ | | ■ | | ■ | | 44 004 | 7 657 |
| Psocidae | Small | Weak | ■ | | | | | ■ | | | | ■ | | | | ■ | | | | ■ | ■ | | | ■ | | | 3 083 | 3 434 |
| Stenopsocidae | Small | Weak | | | | | | | | | | | | | | | ■ | | | | ■ | | | | | | 0 | 1 000 |
| Sminthuridae | Small | Weak | ■ | | | | | | | | | | | | | | | | | | | | | | | | 594 | 0 |
| Hydrodromidae | Small | Weak | | | | | | | | | ■ | | | | | | | | | | | | | | | | 339 | 0 |
| Penthaleidae | Small | Weak | | ■ | | | | | | | | | | | | | | | | | | | | | | | 362 | 0 |
| Pygmephoridae | Small | Weak | | ■ | | | | | | | | | | | | | | | | | | | | | | | 1 628 | 0 |
| Trombidiidae | Small | Weak | | | ■ | | | | | | | | | | | | | | | | | | | | | | 188 | 0 |
