## Supplementary material for "Establishing insect community composition using metabarcoding of soil samples, and preservative ethanol and homogenate from Malaise trap catches: surprising inconsistencies between methods": Table S5

### Table of Contents:


| | |
|---|---|
| **Tables** | Page 1 |
| **Figures** | Page 5 |
| **Summary of the bioinformatic pipelines** | Page 11 |


**Table S1**. Sampling localities with coordinates and vegetation.

| Site | Coordinates | Vegetation |
|---|---|---|
| S1 | 59° 17' 38" N 18° 07' 18" E | Mixed *Betula* and *Picea*. Understory of Blueberry (*Vaccinium,* Ericaceae) and Raspberry *(Rubus,* Rosaceae). |
| S2 | 59° 17' 46" N 18° 09'36" E | *Pinus*. Understory of Blueberry (Vaccinium, Ericaceae). |
| S3 | 59° 17' 42" N 18° 08' 53" E | Forest edge with *Pinus* and *Quercus*. Understory of Blueberry (*Vaccinium*, Ericaceae) and grass (Poaceae). Close to water causes. |

**Table S2**. Number of MOTUs and relative abundance of reads for each arthropod order from all the samples of trap catches combined (ethanol + tissue), from 16S and COI markers.

| | **16S** | | **COI** | |
|---|---|---|---|---|
| **Order** | MOTUs | Reads (%) | MOTUs | Reads (%) |
| Diptera | 209 | 43.68 | 206 | 37.03 |
| Hymenoptera | 25 | 15.64 | 79 | 12.68 |
| Lepidoptera | 29 | 6.36 | 44 | 13.29 |
| Coleoptera | 21 | 5.65 | 33 | 6.42 |
| Hemiptera | 21 | 6.33 | 21 | 5.09 |
| Psocoptera | 6 | 2.21 | 5 | 2.15 |
| Neuroptera | 2 | 0.09 | 2 | 0.19 |
| Trichoptera | 0 | 0.00 | 1 | 0.01 |
| Orthoptera | 1 | 0.13 | 0 | 0.00 |
| Ephemeroptera | 1 | 0.17 | 1 | 0.15 |
| Blattodea | 1 | 0.43 | 0 | 0.00 |
| Symphypleona | 0 | 0.00 | 1 | 0.06 |
| Entomobryomorpha | 1 | 10.32 | 4 | 3.89 |
| Unidentified Collembola | 1 | 1.20 | 0 | 0.00 |
| Araneae | 0 | 0.00 | 6 | 0.91 |
| Trombidiformes | 1 | 0.02 | 6 | 0.45 |
| Mesostigmata | 0 | 0.00 | 3 | 0.20 |
| Sarcoptiformes | 0 | 0.00 | 2 | 0.51 |
| Opiliones | 1 | 0.30 | 1 | 0.14 |
| Other Metazoa | 0 | 0.00 | 0 | 0.00 |
| Other kingdoms | 0 | 0.00 | 5 | 0.31 |
| Unknown | 17 | 7.47 | 5 | 16.52 |

**Table S3**. Number of MOTUs and relative abundance of reads for each arthropod order from all the soil samples combined (humus + leaf litter), from 16S and COI markers.

| | **16S** | | **COI** | |
|---|---|---|---|---|
| **Order** | MOTUs | Reads (%) | MOTUs | Reads (%) |
| Diptera | 34 | 15.68 | 0 | 0.00 |
| Hymenoptera | 1 | 0.06 | 0 | 0.00 |
| Coleoptera | 11 | 12.34 | 2 | 0.06 |
| Hemiptera | 9 | 3.74 | 2 | 3.24 |
| Psocoptera | 3 | 0.25 | 0 | 0.00 |
| Orthoptera | 0 | 0.00 | 1 | 0.06 |
| Symphypleona | 1 | 0.05 | 1 | 0.01 |
| Entomobryomorpha | 10 | 12.66 | 0 | 0.00 |
| Neelipleona | 2 | 0.22 | 0 | 0.00 |
| Poduromorpha | 8 | 12.91 | 3 | 0.03 |
| Unidentified Collembola | 23 | 22.74 | 1 | 0.03 |
| Araneae | 0 | 0.00 | 2 | 0.17 |
| Trombidiformes | 1 | 0.19 | 1 | 0.02 |
| Opiliones | 1 | 0.01 | 0 | 0.00 |
| Isopoda | 1 | 0.01 | 0 | 0.00 |
| Julida | 1 | 2.20 | 1 | 0.03 |
| Lithobiomorpha | 1 | 0.22 | 0 | 0.00 |
| Other Metazoa | 2 | 2.39 | 6 | 0.24 |
| Other kingdoms | 0 | 0.00 | 32 | 53.48 |
| Unknown | 13 | 14.34 | 36 | 42.65 |

**Table S4.** Families recovered exclusively by 16S or COI markers from each of the trap samples (*i.e.* bulk tissue or preservative fluid), or detected by both markers.

| Sample | 16S | COI | Both |
|---|---|---|---|
| S1T1 | 9 | 15 | 22 |
| S1T2 | 5 | 16 | 17 |
| S1T3 | 5 | 7 | 20 |
| S1T4 | 4 | 12 | 13 |
| S2T1 | 6 | 14 | 24 |
| S2T2 | 12 | 8 | 18 |
| S2T3 | 8 | 10 | 17 |
| S2T4 | 3 | 12 | 12 |
| S3T1 | 6 | 11 | 21 |
| S3T2 | 13 | 13 | 19 |
| S3T3 | 5 | 4 | 16 |
| S3T4 | 2 | 14 | 11 |

**Table S5**. List of families recovered exclusively from ethanol or tissue, or both from ethanol and tissue, ordered by their degree of sclerotization and size (hard and large at the top, small and weak at the bottom). A blue square indicates that representatives of the family were detected in ethanol, and a red square that they were detected in tissue. The number of reads of each sample type is indicated in the rightmost two columns. [EXCEL FILE]

**MOTU table from 16S** [CSV FILE]

**MOTU table from COI** [CSV FILE]

**Figure S1**. Number of clusters (MOTUs) generated by SWARM at different nucleotide distance thresholds for the 16S–'short', 16S–'long' and COI. When the number of MOTUs stabilizes it means no more oversplitting of species is occurring.

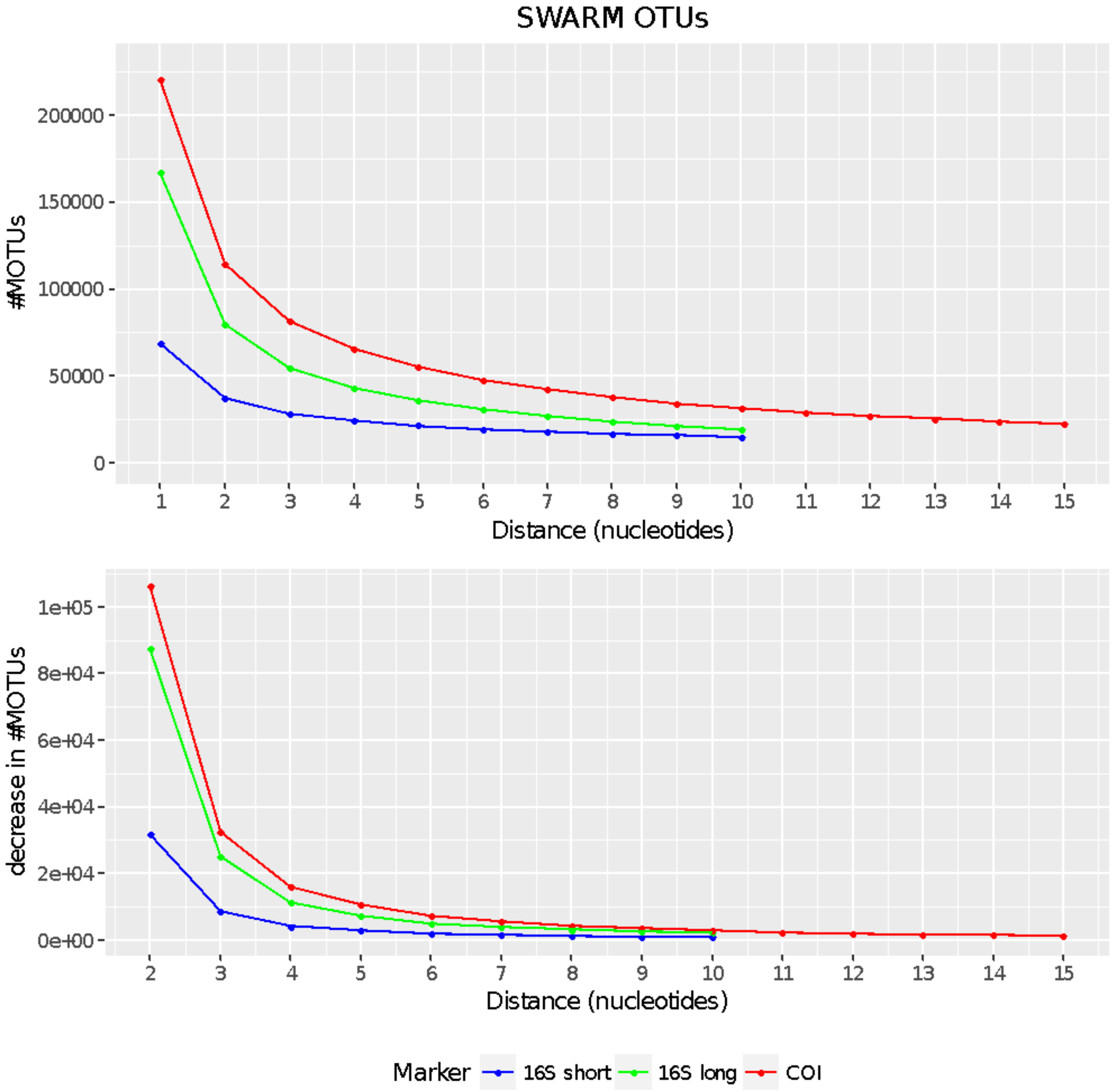

**Figure S2**. Proportion of reads corresponding to MOTUs identified as Arthropoda and non-Arthropoda by 16S (upper panel) and COI (lower panel) from each sample. MOTUs unidentified at the phylum level are included within the non-Arthropoda.

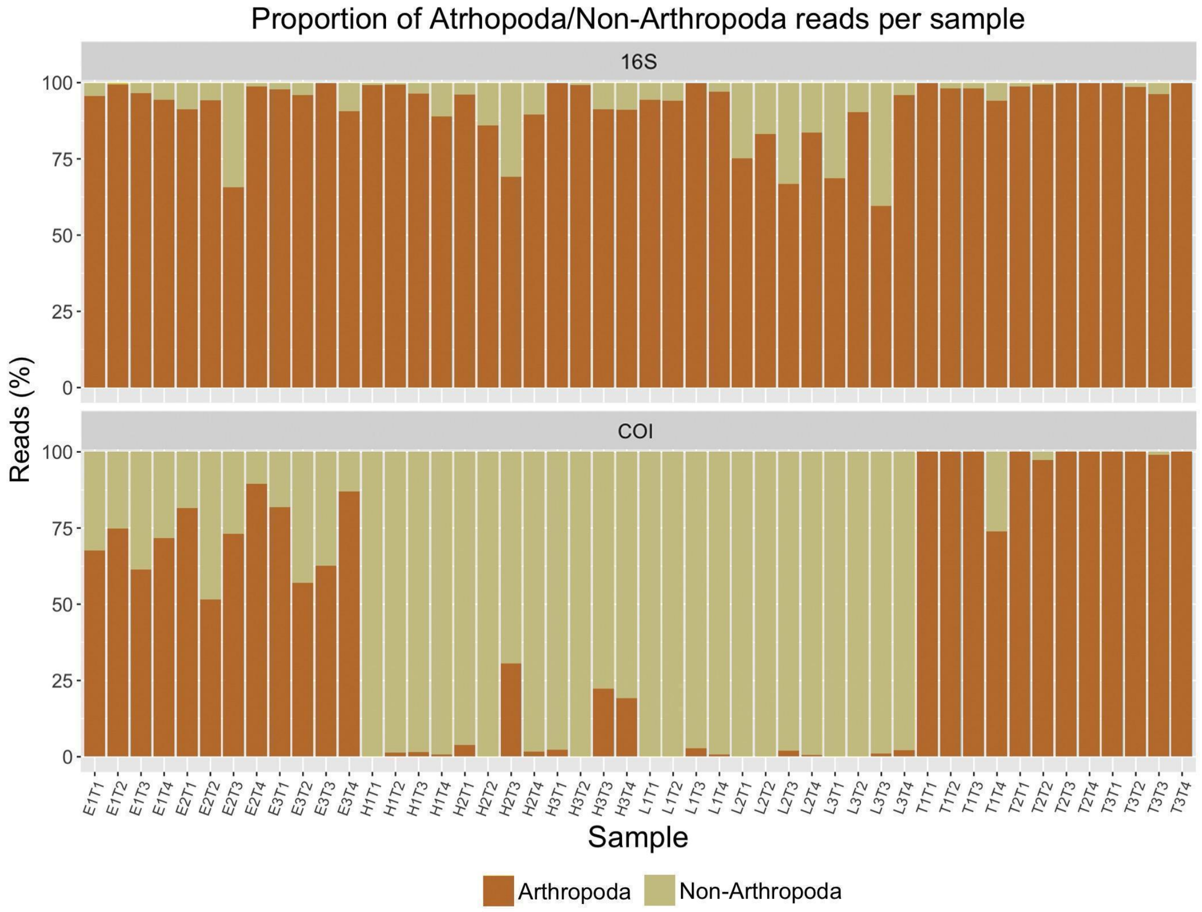

**Figure S3**. Rarefaction curves from the trap samples (ethanol and tissue). **A)** Ethanol 16S, **B)** Tissue 16S, **C)** Ethanol COI, **D)** Tissue COI. The number of reads of the sample with the least reads is indicated with a dashed line.

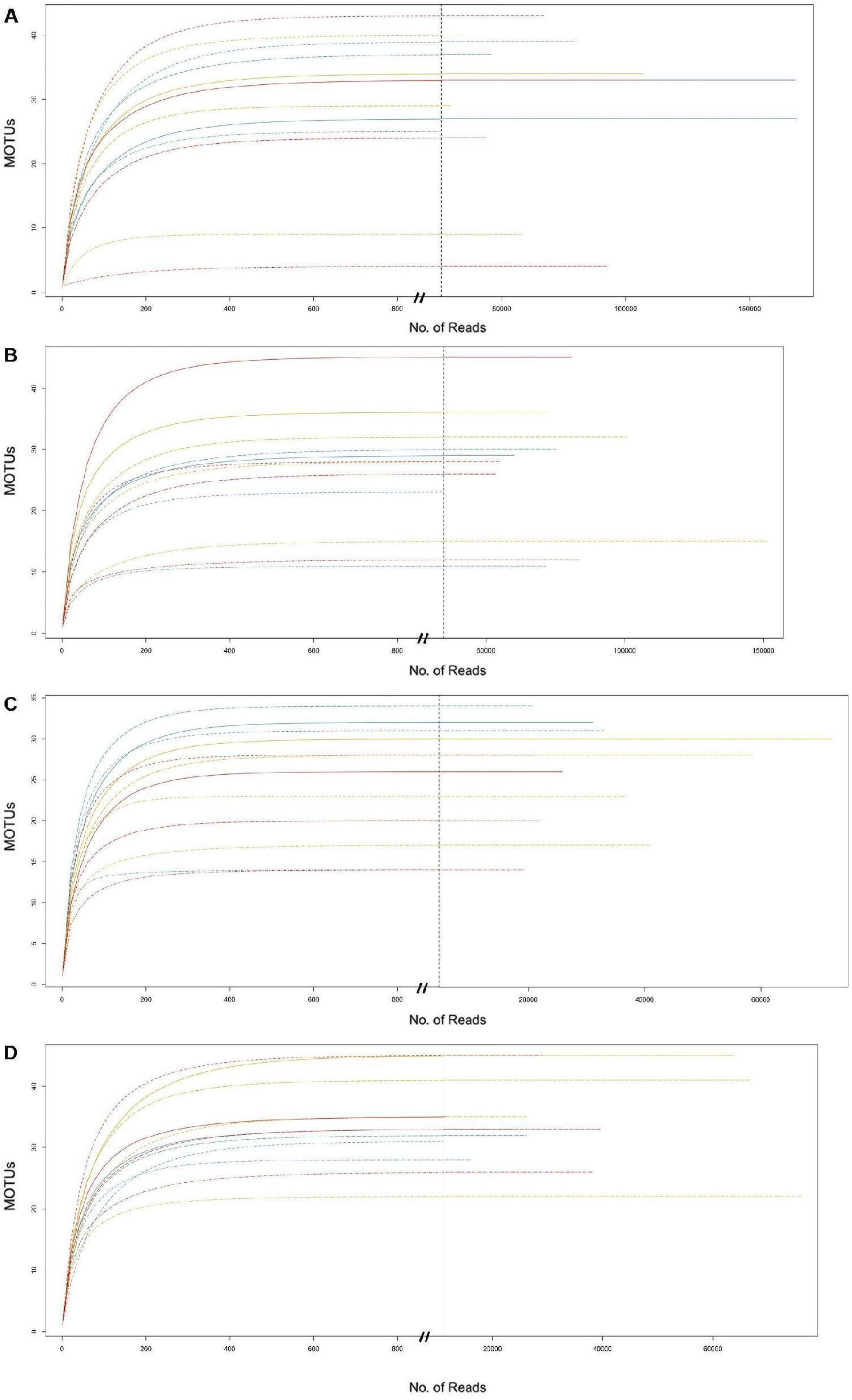

**Figure S4**. Rarefaction curves from the trap samples (ethanol and tissue). **A)** Humus 16S, **B)** Leaf litter 16S, **C)** Humus COI, **D)** Leaf litter COI. The number of reads of the sample with the least reads (different from zero) is indicated with a dashed line. A red asterisk (*****) indicates samples that did not reach the stationary phase (this occurred for sample H2T2, the number of reads of which was 72).

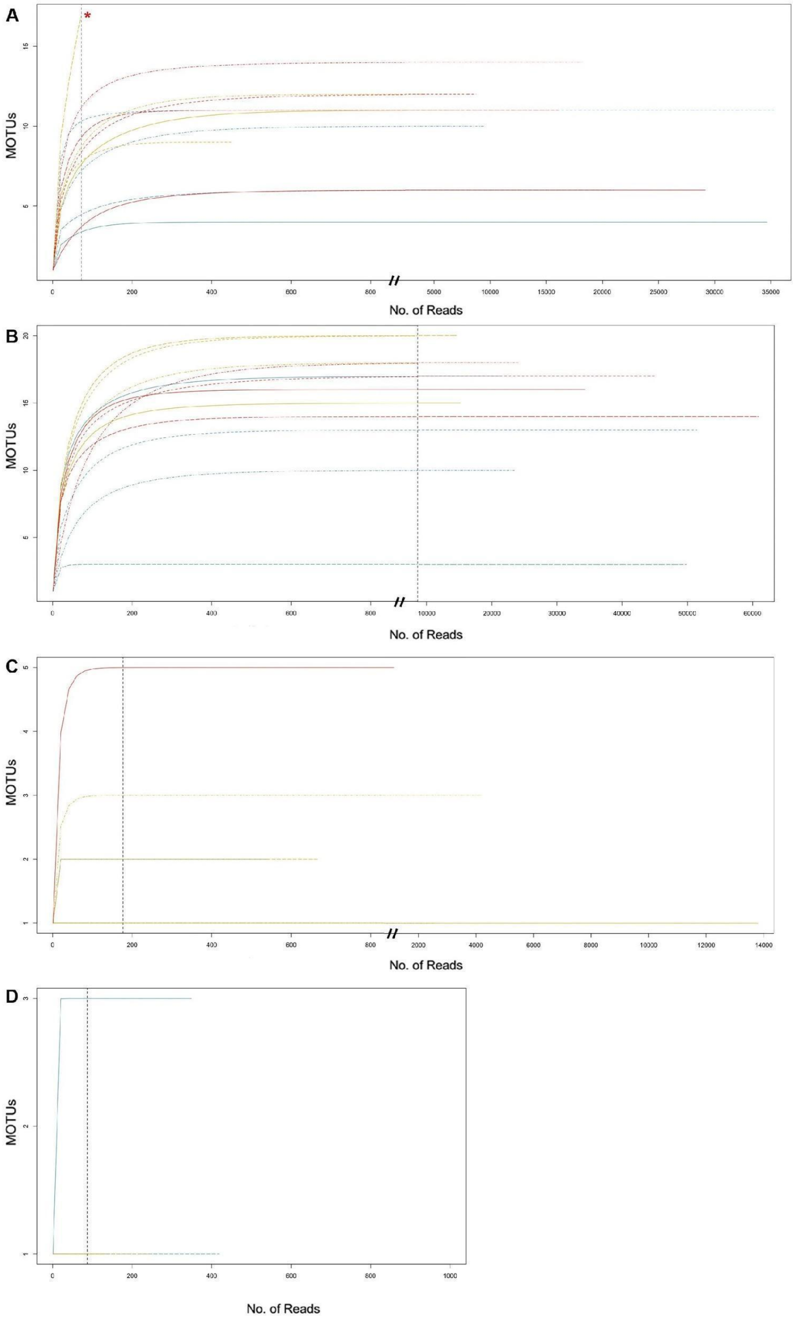

**Figure S5**. Number of MOTUs recovered exclusively from ethanol or tissue, or from both ethanol and tissue, as split by sample.

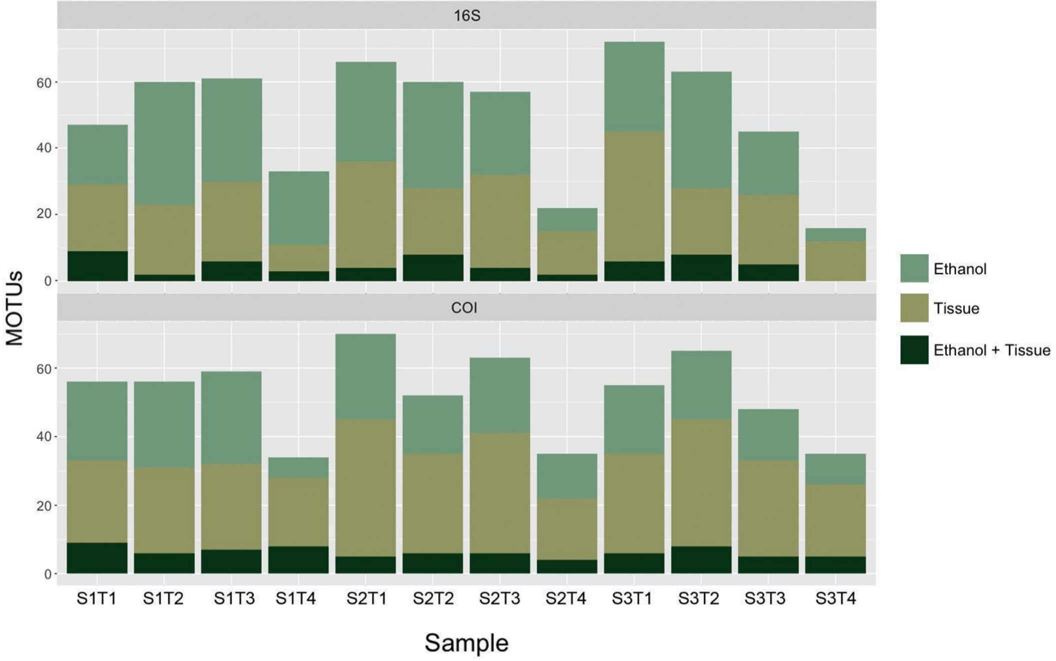


**Figure S6**. Number of MOTUs recovered exclusively from humus or leaf litter, or both from ethanol and tissue, as split by sample.

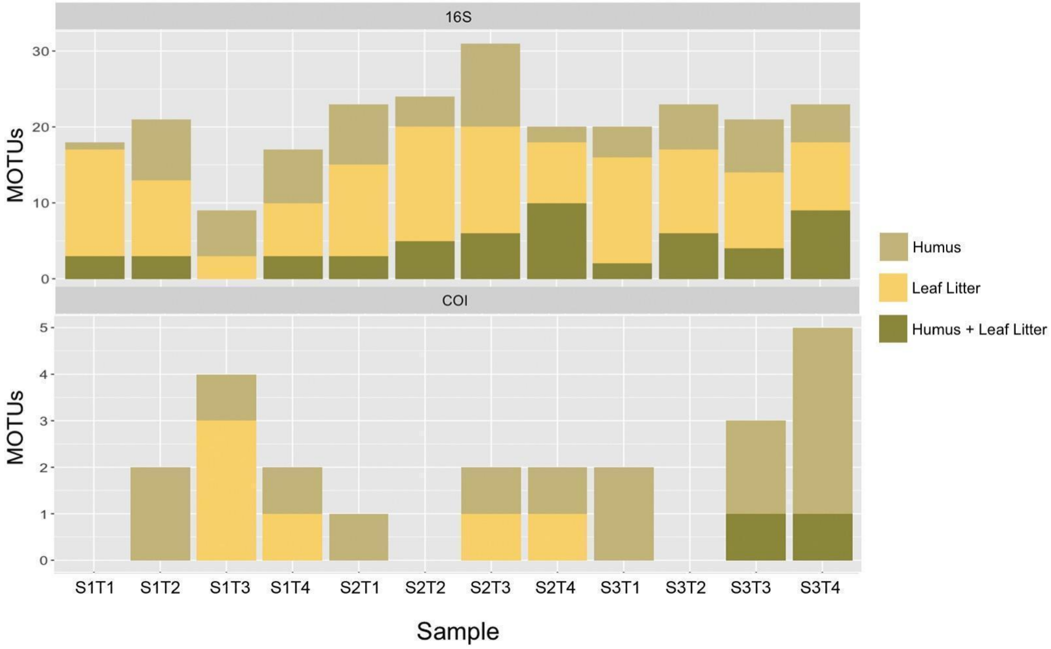

**Figure S7**. Number of MOTUs recovered exclusively from soil (humus and leaf litter) or trap substrates (ethanol and tissue), or both from soil and traps, as split by sample.

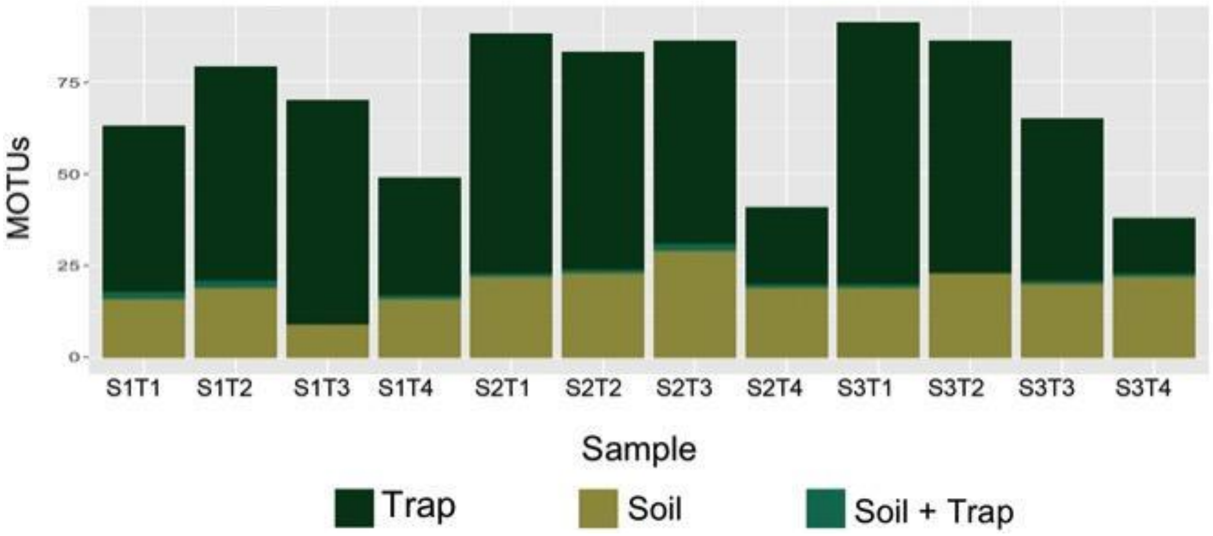

### Summary of the bioinformatic pipelines

| Process | Software | Options | Source |
|---|---|---|---|
| Quality assessment of reads and trimming | FastQC<br>obicut | -<br>- | -<br>OBITools |
| Paired-end merging | illuminapairedend | Discarded score<30.00 | OBITools |
| Demultiplexing | ngsfilter | --fasta-output -u unidentified | OBITools |
| Length selection | obigrep | -p 'seq_length>310' -p 'seq_length<330' (COI) / -p 'seq_length>315' -p 'seq_length<375' (16S-'long') / -p 'seq_length>255' -p 'seq_length<305' (16S-'short')<br>-s '^[ACGT]+$' | OBITools |
| Dereplication | obiuniq | -m sample | OBITools |
| Reads abundance counting | obitab | | OBITools |
| Chimera removal | uchime_denovo | --sizeout<br>--nonchimeras *nonchimeras.fasta* | VSEARCH |
| Clustering | swarm | -d 9 (COI) / -d 5 (16S-'long') / -d 4 (16S-'short')<br>- z<br>-o *outputfile*<br>-w *seeds.fasta* | SWARM |
| MOTUs abundance counting | owi_reciunt_swarm | - | Metabarpark |
| Matchlist seed sequences | usearch_global | --db *seeds.fasta*<br>--self<br>--id .84<br>--userout *matchlist.txt*<br>--userfields query+target+id<br>--maxacepts 0<br>--query_cov .9<br>--maxhits 10 | VSEARCH |
| Co-ocurrence curation | lulu | - | LULU |
| Taxonomic identification | ecotag<br>owi_add_taxonomy | - | OBITools<br>Metabarpark |

| | | | |
|---|---|---|---|
| MOTUs table | owi_combine | - | Metabarpark |
| Low absolute and relative (per sample) abundance correction | finalMOTUs_curation | 0.004<br>10 | Metagusano |

Sources

- OBITools: https://pythonhosted.org/OBITools/welcome.html
- VSEARCH: https://github.com/torognes/vsearch
- SWARM: https://github.com/torognes/swarm
- Metabarpark: https://github.com/metabarpark/R_scripts_metabarpark
- -LULU: https://github.com/tobiasgf/lulu
- Metagusano: https://github.com/metagusano/metabarcoding_scripts
